# Variability in foraging ranges of snow petrels and implications for breeding distribution and use of stomach-oil deposits as proxies for paleoclimate

**DOI:** 10.1101/2025.06.06.658237

**Authors:** Ewan D. Wakefield, Erin L. McClymont, Sébastien Descamps, W. James Grecian, Eleanor M. Honan, Anna S. Rix, Henri Robert, Vegard Sandøy Bråthen, Richard A. Phillips

## Abstract

**Background:** Breeding pelagic seabirds feed over wide areas. Their diet and space use can therefore provide proxies for otherwise be difficult to observe environmental conditions but interpretation of these proxies requires knowledge of foraging range. Antarctic sea ice regulates global climate, but a paucity of data on its past extent causes uncertainty in climate reconstruction and prediction. Deposits of stomach oils, produced defensively by snow petrels *Pagodroma nivea* (near-obligate sea-ice foragers), reflect sea ice conditions, sometimes over tens of millennia, but spatial interpretation of these biological archives is hampered by lack of data on foraging range, and how this varies seasonally, among colonies and with sex.

**Methods:** We tracked 34 and 60 snow petrels using light-based geolocators and GPS loggers, respectively to estimate foraging ranges at three colonies located 180-200 km inland in Dronning Maud Land. We tested whether foraging latitude is associated with ice edge latitude, estimated via satellite remote sensing. We then projected potential foraging ranges for all known colonies on the study area to reexamine assumptions made in paleoclimate studies.

**Results:** During most breeding stages, and across breeding seasons, core home ranges were centred approximately 2° south of the outer sea-ice edge and tracked this habitat as it receded during the spring melt. Females were approximately 7% lighter than males but foraged at similar distances and in similar areas. Foraging ranges differed little between colonies but substantially between breeding stages. For example, median range was ∼1,400 km during the pre-laying exodus vs. ∼550 km during brood-guard.

**Conclusions:** Snow petrel stomach-oil deposits potentially integrate environmental conditions over greater and more seasonally variable areas than previously assumed, probably with a bias towards conditions in the outer Marginal Ice Zone during the early summer when stomach oil deposition due to nest competition is likely greatest. Projected potential home ranges of snow petrel colonies in the western Weddell Sea suggest that breeding is limited by access to foraging habitat, such as coastal polynyas. We hypothesise that as sea ice fluctuated over previous glacial-interglacial cycles, this also regulated breeding distribution across the region.

## 1. Background

Knowledge of animal movements and distributions is fundamental to ecology and wildlife management. Breeding seabirds are central-place foragers, alternating between foraging at sea and courting, defending their nests, incubating their egg(s) and brooding, guarding or feeding their offspring. These duties place energetic and temporal limits on foraging range [1–3], defined here as the distance that a seabird can travel from the nest to gather food while breeding successfully. If foraging range is known, seabird diet and behaviour can be used to infer environmental conditions within a known area at sea. A promising new example is the analysis of stomach-oil deposits from snow petrels *Pagodroma nivea* to infer past climatic and biological conditions around Antarctica [4–6].

Snow petrels breed in the austral summer in cavities in or among snow- and ice-free rocks [7, 8], and forage mainly in association with sea ice [9–11]. Like other Procellariformes, their proventriculus is adapted to separate and retain the energy-rich lipid fraction of their prey both for their own sustenance, and to feed to their chick [12, 13]. They also spit stomach oil at conspecifics during agonistic competition for nest sites and as a defence against their main predator, the south polar skua *Stercorarius maccormicki* [8, 14]. Over periods sometimes >50,000 years, stomach oil builds up around nest sites in waxy, stratified accretions up to tens of cm thick [15–17]. The accumulation rates and biochemistry of these deposits reflect temporal variation in nest occupation and diet, providing proxies for past sea-ice conditions and the timing of ice-sheet advance or retreat in the nesting locality [4–6, 18, 19]. Potentially, this could fill an information gap on sea-ice extent in the pre-satellite era, which is a major source of uncertainty in climate models [20, 21], particularly south of the Antarctic Polar Front [22–24]. However, the utility of stomach-oil deposits as climate proxies is currently limited by lack of data on snow petrel foraging ranges. This also hampers understanding of the effects of the availability and accessibility of sea-ice foraging habitats on their past, current and future nesting distribution [25, 26].

Potential foraging range (i.e., the distance reachable in the absence of extrinsic factors) is ultimately limited by morphological and physiological constraints on locomotion and body maintenance, modulated by temporal and energetic demands that vary across the breeding cycle [1, 27], and with sex, especially among size-dimorphic species [28, 29]. Realised foraging ranges are the product of further modulation by extrinsic effects: prey and suitable foraging habitats are patchily distributed [30]; movement constraints may necessitate indirect commuting routes [27]; and intraspecific competition cause spatial segregation among adjacent colonies and increase foraging range [31, 32].

Snow petrels usually forage in areas of intermediate (∼30 – 60 %) sea-ice concentration, i.e. those typical of the Marginal Ice Zone (MIZ) [9–11, 33]. They breed from October to March [8], and Antarctic sea ice is at its maximum and minimum extents in September and February, respectively [34]. Hence, if snow petrels track the MIZ over the spring melt, they should forage further south, with decreasing foraging range, as the breeding season progresses and the MIZ retreats towards the coast [35–37]. Breeding distribution is expected to be limited by distance suitable foraging habitat, such as the MIZ [38, 39]. Ship-based surveys suggest that the highest densities during breeding are within ∼400 km of colonies [10, 37], but recent tracking indicates foraging ranges of 100s to 1000s of km [11, 40]. Variation in foraging ranges over the breeding cycle, the effects of taking indirect routes due to the constraints of topography, wind, etc. [27, 41], and whether there is spatial segregation among colonies with potentially overlapping foraging areas are all unknown. Similarly, little is known the effect of sex on foraging range other than one study which showed that despite snow petrels being sexually size dimorphic, incubating males and females had similar foraging ranges [11].

The Weddell Sea sector of the Southern Ocean (Fig. 1) is important in terms of sea-ice-climate linkages [42]. Snow petrels breed in its hinterland throughout Dronning Maud Land (DML), and west to the Theron Mountains and Shackleton Range [7]. Stomach-oil deposits from the region have been analysed to infer past occupation history [16, 17], variation in sea-ice conditions since the last glacial [5, 6, 43] and ice-sheet thinning history [18, 44], but foraging ranges of snow petrels breeding in the region are unknown. Recent loss of seasonal sea ice has been particularly pronounced in the Weddell Sea sector [45, 46] – a trend predicted to continue under anthropogenic climate change [21, 47]. Development of marine protection measures is currently hampered by a lack of data, especially west of the prime meridian [48].

**Fig. 1.**
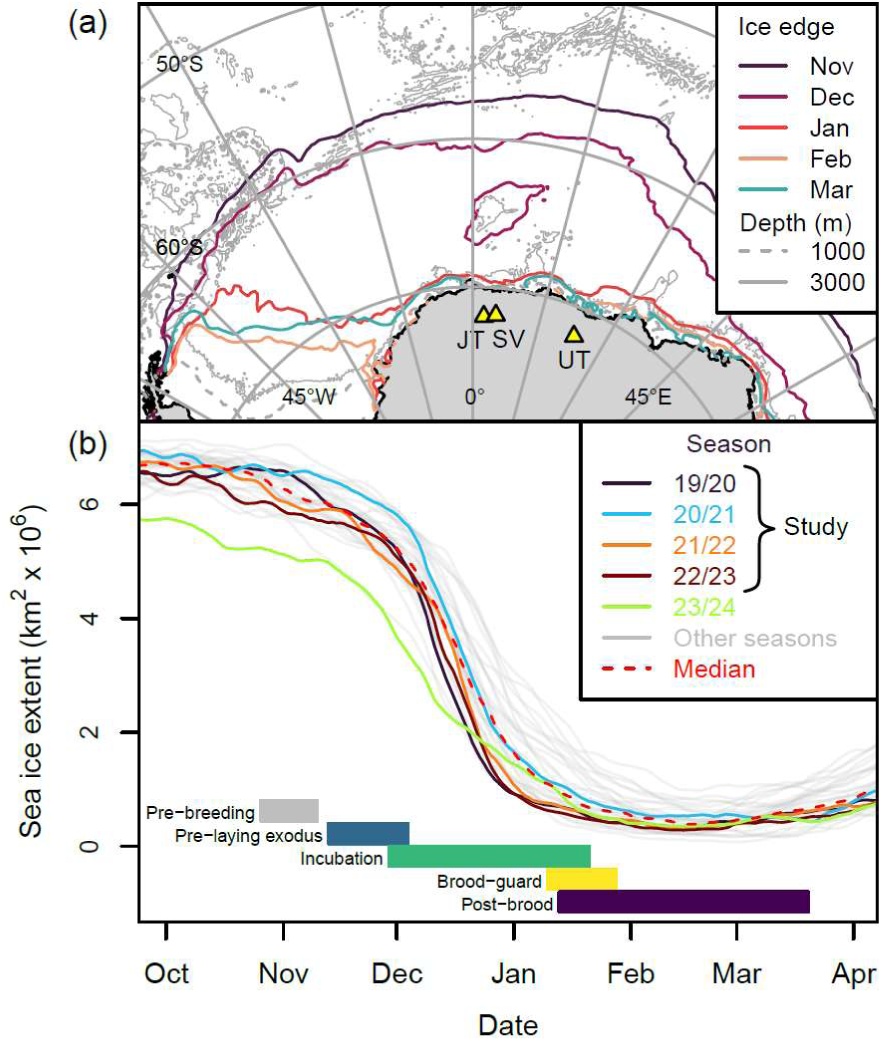
(a) Study area showing the 30-year monthly median ice edge during the snow petrel breeding season, study colonies (yellow triangles), and places mentioned in the text (JT Jutulsessen; MR Maud Rise (location of seasonal polynya); TT Troll Ice Tongue; SV Svarthamaren; UT Utsteinen). (b) Evolution of sea-ice cover in the sector 040°W to 040°E during each of the 30 snow petrel breeding seasons from 1994/95 to 2023/24, in relation to the snow petrel breeding cycle.

## 2. Methods

### 2.1 Aims

Here, we aim to estimate the foraging ranges and distributions of snow petrels and test whether they vary with breeding stage, sex and colony. To do so, we tracked birds from three colonies in DML. We illustrate the application of our results for the interpretation of climate proxies derived from stomach-oil deposits, and the relationship between sea-ice conditions and breeding-site occupation by projecting potential foraging ranges of snow petrels from colonies throughout the Weddell Sea region.

### 2.2 Study system and data collection

The coastline of DML (020°W-045°E) principally comprises ice shelves which occlude the underlying bedrock [48], and the snow petrels nest 100 - 290 km inland on a discontinuous chain of nunataks, running parallel to the coast [7, 49]. Two forms that differ in size - greater and lesser - have been recognised [50], but these intergrade and their taxonomic status is unresolved [51]. Birds breeding in DML are all considered to be of the lesser form [8]. We tracked breeding snow petrels from three colonies located 203, 183 and 190 km from the coast (Fig. 1): Jutulsessen (72°02.1’S, 002°28.2’E; ∼3000 pairs); Svarthamaren (71°53.4’S, 005°09.6’E; ∼2000 pairs); and Utsteinen, (∼200 pairs) [52, 53, authors’ unpub. data]. Jutulsessen and Svarthamaren are Important Bird Areas and the latter is an Antarctic Specially Protected Area (ASPA) [54].

The breeding schedule of snow petrels (Fig. 1) is similar across Antarctic continental breeding sites [8, 40, 55, 56]. During the spring melt, sea ice off DML recedes southward in the east of the region and south-westwards in the west (Fig. 1a). Open water often occurs over the Maud Rise before the outer ice edge has receded to this feature and breakup is most rapid during December. Minimum sea-ice extent is reached in February (Fig. 1b), when open water occurs along much of the DML coast, but sea ice persists in the western Weddell Sea year-round (Fig. 1a).

Due to logistical constraints, fieldwork was feasible only between late November and late February, i.e. incubation through to mid chick-rearing. Combining GPS loggers (late breeding season) and light-based Global Location Sensors (GLS) allowed bird movements to be tracked throughout the entire breeding period. In our study system, light-based geolocation is only feasible from early October to mid-November and early February to mid-March (Fig. S1). The location accuracies of GPS and GLS tracking are approximately ± 20 m (PathTrack, unpub. data) and ± 200 km, respectively [57–59].

We caught breeding adults at the nest, weighed them (±2.5 g), removed three contour feathers for molecular sexing (see Supplementary Methods) and attached either a GPS or GLS logger. We attached NanoFix GEO remote-download GPS loggers (Pathtrack Ltd., Otley, UK; 3.7 ± 0.3 g (range 3.1-4.1 g); 43 x 15 x 9.5 mm, plus 50 mm whip antenna; most fitted with a solar panel) to the base of the middle two rectrices using Tesa tape and C65-SUPER combined GLS-immersion loggers (Migrate Technology, Cambridge, UK; 1 g; 14 × 8 × 6 mm) to the left tarsus using a plastic ring. We deployed GPS loggers at Svarthamaren during incubation, brood-guard and post-brood (2022/23) and at Jutulsessen during post-brood (2024); and GLS loggers at Svarthamaren and Utsteinen between 2019 and 2023 (Table 1). Mean masses of GPS and GLS loggers plus attachment materials were 2.0 ± 0.3 % (range 1.3-2.7) and 0.8 ± 0.1 % (range 0.5-1.0) of body masses, respectively. Total handling time was 7 ± 2 minutes (range 4-13).

**Table 1.** Numbers of tracking devices deployed on snow petrels at colonies in Dronning Maud Land and from which data were recovered, 2019/20-2023/24. Season Colony.

| Season | Colony | N deployed, recovered (female, male, unknown) |  |
| --- | --- | --- | --- |
|  |  | GLS | GPS |
| 2019/20 | Svarthamaren | 20, 0 |  |
| 2021/22 | Svarthamaren | 6, 0 |  |
|  | Utsteinen | 12, 0 |  |
| 2022/23 | Svarthamaren | 15, 5 (0,3,2) | 44, 36 (11,25,0) |
|  | Utsteinen | 0, 7 (4,3,0) |  |
| 2023/24 | Svarthamaren | 0, 22 (2,8,12) |  |
|  | Jutulsessen | 0, 0 | 31, 24 (11,13,0) |

GLS loggers recorded maximum light intensity every five minutes and wet-dry state either as the number of samples within ten-minute blocks when the logger was immersed, based on a test every 30 s, or the times of state changes between wet and dry sampled every 6 s. GPS loggers acquire a fix every 30 minutes when the battery was fully charged or at longer intervals when it was depleted. If possible, GPS loggers deployed during incubation were recovered that season. Those deployed during chick-rearing were not recovered but continued to transmit data to base stations until the tags failed or were lost. Snow petrels moult their rectrices at the end of the breeding season, or earlier if breeding fails [8], so loggers not recovered would have been shed by April. We recovered GLS loggers after 1-4 y, with recovery attempts in all seasons except 2020/21.

All sea-ice metrics are based on analysis of sea-ice concentration (SIC) data downloaded from the National Snow and Ice Data Centre [60] on a 25 km regular grid [61]. To define the ice edge, we followed the commonly applied operational assumption that this corresponded to the 15% SIC contour.

### 2.3 GLS data processing and colony attendance

Following [62], we used the light level data to obtain twice daily locations estimates (see Supplementary Methods). In brief, we used the threshold method to estimate twilight times, and calculated latitude from the inter-twilight interval and longitude from the time of noon or midnight. We did not attempt to make location estimates during equinoctial periods or during bouts of daylight exceeding 24h (i.e. when birds were at latitudes of >°S during midsummer). We filtered both twilights and positions to reduce location errors.

Prior to analysis, we reduced activity data from all loggers to the proportion of each ten-minute period immersed. We then defined putative bouts in the colony as those during which loggers were immersed for <95% of each 24 h period and bouts on the nest as those in which loggers were shaded (based on the light data) throughout most of the day. We assumed that birds still carrying out nest shifts after January 1^st^ hatched their chick successfully that season and considered all other birds to have been breeding up to the end of their last recorded incubation shift. For subsequent analyses, we retained only locations at sea, defined using the High Resolution Vector Polygons of the Antarctic Coastline (7.6) dataset [63].

### 2.4 GPS data processing and behavioural classification

Prior to analysis, we split GPS tracks into foraging trips and interpolated locations to 30-minute intervals using a correlated random walk model (Supplementary Methods). The higher resolution of GPS than GLS data also allowed us to exclude bouts of non-foraging behaviours prior to estimation of foraging range, as these potentially occur non-uniformly with respect to distance from the colony. To do so we followed Wakefield et al. [35], using Hidden Markov Models to classify movement based on step length and turning angle as either travelling (step length high, turning angle concentrated), foraging (step length intermediate, turning angle dispersed) or resting (step length low, turning angle very dispersed) [64, 65]. In brief, we implemented models in moveHMM package [66]. We fitted HMMs 25 times, randomly drawing starting parameters from within plausible ranges based on previous studies [35, 67] (Table S1). Using the model with the highest likelihood [68], we predicted the most likely sequence of states using the Viterbi algorithm, retaining only the putative foraging locations for subsequent analyses.

### 2.5 Estimation of space use and comparisons among groups

To summarise space use, we used the adehabitatHR package [69] to estimate utilisation distributions (UDs) via kernel analysis, specifying a smoothing parameter of 100 km for GLS and 20 km for GPS data, respectively. For the latter, we first estimated UDs for each bird/stage/season and then averaged these to obtain UDs for stages within seasons. For GLS-tracked birds (two-locations/day), there were insufficient data to first estimate bird-level UDs, so we aggregated locations across birds prior to estimating UDs for stages within seasons. We define contours containing the first 50 and 75% percent of cumulative utilisation, as core home ranges and home ranges, respectively.

We used two approaches to compare space use overlap among groups. For both GLS and GPS-tracked groups, we calculated the Home Range Overlap Index, *HROI* [31], defined here as the mean of the proportion of each group’s core home range intersected that of the other group’s home range:

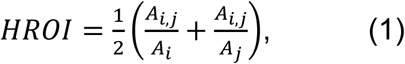

where *A*_*i*_ and *A*_*j*_ are the areas of group *i* and *j*’s core home ranges and *A*_*i,j*_ is the area of their intersection (cf. Fieberg and Kochanny’s [70] *HR*). This ranges from 0 (no overlap) to 1 (complete overlap). Secondly, we quantified similarity among the UDs of GPS-tracked groups using Bhattacharyya’s affinity [70]:

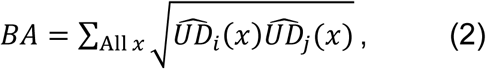

where 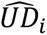 and 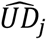 are the observed UDs of birds from groups *i* and *j* on a grid comprising cells *x*. BA also ranges from 0 (complete dissimilarity) to 1 (identical). We tested the null hypothesis that observed UDs did not differ between groups (male vs. female or colony 1 vs. colony 2) by randomly shuffling bird identities across the two groups without replacement and recalculating BA 1000 times, then calculating the *p* value as the proportion of null BA values that were less than the observed value, considering this a one-sided test [11, 71].

### 2.6 Foraging range estimation, comparison, and projection

When commuting between the colony and the coast, birds followed beeline tracks modified by wind drift similar to those of sympatric Antarctic petrels *Thalassoica antarctica* described by Tarroux et al. [72] (see Section 4.2). Once at sea, snow petrels largely avoided crossing land or ice shelves, even when travelling to destinations that could have been reached more directly over land. Moreover, presumably due to prevailing easterly winds, they crossed the coast further west on the outward leg than on the inward leg of most foraging trips (see Results). We therefore used biological distance – i.e., distance via the shortest paths conforming to these constraints [41], to quantify foraging range (see Supplementary Methods). For each individual, we then quantified foraging range as the median (*d*_50_), 95^th^ percentile (*d*_95_) and maximum (*d*_*max*_) biological distance of at-sea locations within each breeding stage. For GPS-tracked birds, we use the subscript *f* to denote statistics that summarise *d* across foraging locations (e.g., *d*_*f*,50_, etc.), whereas for GLS-tracked birds we use the subscript *u*, denoting all forms of utilisation. We defined the mean heading during overland commutes between the colony and the first location at sea as *θ*_*out*_, and between the last location at sea and the colony as *θ*_*in*_.

Tests for differences in foraging range among groups consistently led to the same conclusions whether they were carried out on *d*_*u*,50_ and *d*_*f*,50_ or *d*_*u*,95_ and *d*_*f*,95_ (only the former results are presented for brevity). For simple two-sample or paired data, we used *t* or Wilcoxon tests, checking equality of variance and normality using *F* and Shapiro-Wilk tests. We estimated repeatability *R* [73] in foraging range using the rptR package [74]. We used Generalised Linear Models (GLMs) to model median foraging range derived from GLS data (one observation per individual) and mixed-effects GLMs (GLMMs), fitted in the lme4 package [75], with bird-level random intercepts and inverse Gaussian distributed errors with an inverse link function, to model foraging ranges from the GPS data (multiple observations per individual), which were positively skewed. We scaled the response by dividing by standard deviation prior to model fitting. We checked conformity to model assumptions using Q-Q and residual plots and we carried out multiple comparisons between breeding stages using the package emmeans [76]. Using our results (Table 2), we projected the potential home ranges of birds foraging from all known breeding locations in the study area [7, 77].

**Table 2.**
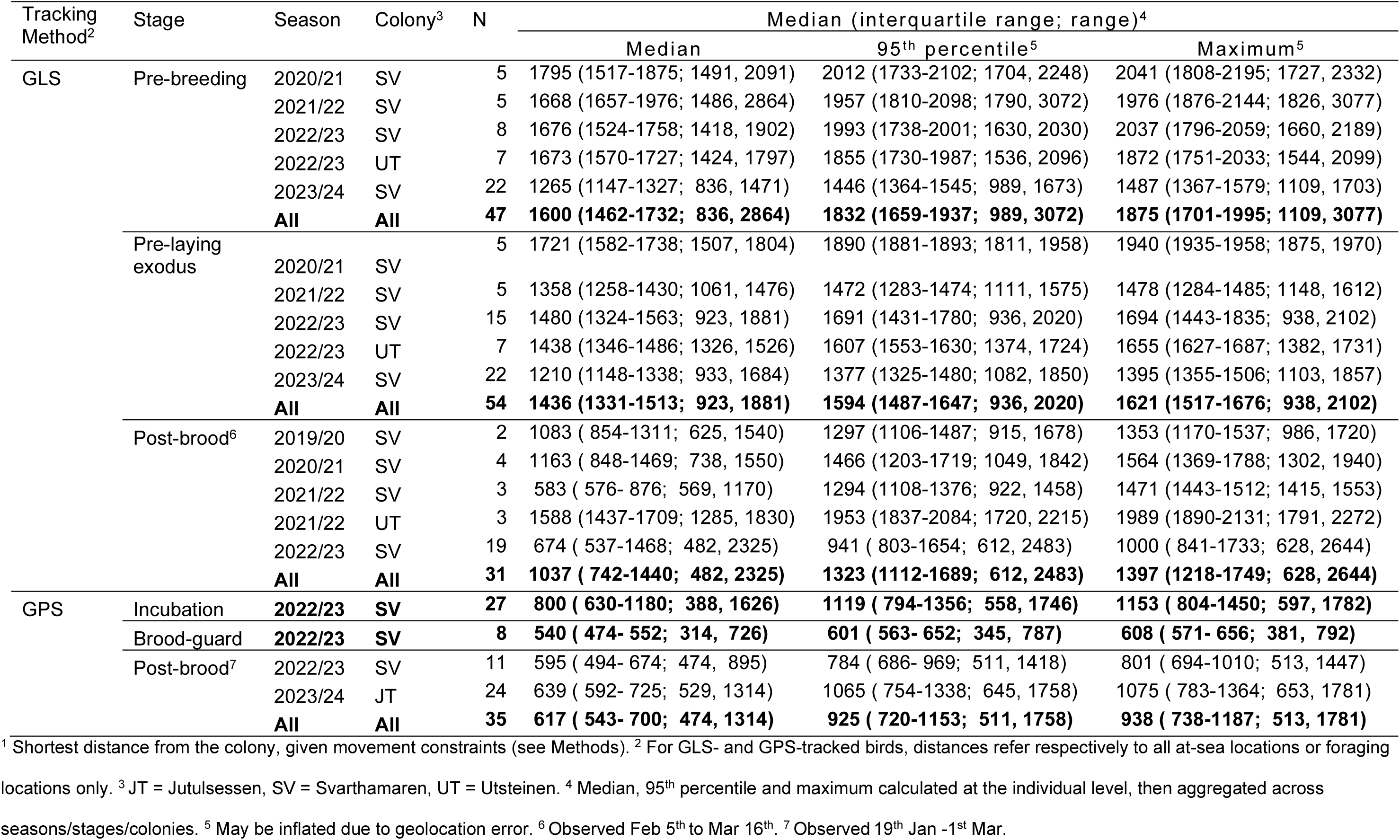
Realised foraging ranges^1^ (km) of snow petrels tracked from three colonies in Dronning Maud in 2020/21-2023/24.

| Tracking Method<sup>2</sup> | Stage | Season | Colony<sup>3</sup> | N | Median (interquartile range; range)<sup>4</sup> |  |  |
| --- | --- | --- | --- | --- | --- | --- | --- |
|  |  |  |  |  | Median | 95<sup>th</sup> percentile<sup>5</sup> | Maximum<sup>5</sup> |
| GLS | Pre-breeding | 2020/21 | SV | 5 | 1795 (1517-1875; 1491, 2091) | 2012 (1733-2102; 1704, 2248) | 2041 (1808-2195; 1727, 2332) |
|  |  | 2021/22 | SV | 5 | 1668 (1657-1976; 1486, 2864) | 1957 (1810-2098; 1790, 3072) | 1976 (1876-2144; 1826, 3077) |
|  |  | 2022/23 | SV | 8 | 1676 (1524-1758; 1418, 1902) | 1993 (1738-2001; 1630, 2030) | 2037 (1796-2059; 1660, 2189) |
|  |  | 2022/23 | UT | 7 | 1673 (1570-1727; 1424, 1797) | 1855 (1730-1987; 1536, 2096) | 1872 (1751-2033; 1544, 2099) |
|  |  | 2023/24 | SV | 22 | 1265 (1147-1327; 836, 1471) | 1446 (1364-1545; 989, 1673) | 1487 (1367-1579; 1109, 1703) |
|  |  | **All** | **All** | **47** | **1600 (1462-1732; 836, 2864)** | **1832 (1659-1937; 989, 3072)** | **1875 (1701-1995; 1109, 3077)** |
|  | Pre-laying exodus |  |  | 5 | 1721 (1582-1738; 1507, 1804) | 1890 (1881-1893; 1811, 1958) | 1940 (1935-1958; 1875, 1970) |
|  |  | 2020/21 | SV |  |  |  |  |
|  |  | 2021/22 | SV | 5 | 1358 (1258-1430; 1061, 1476) | 1472 (1283-1474; 1111, 1575) | 1478 (1284-1485; 1148, 1612) |
|  |  | 2022/23 | SV | 15 | 1480 (1324-1563; 923, 1881) | 1691 (1431-1780; 936, 2020) | 1694 (1443-1835; 938, 2102) |
|  |  | 2022/23 | UT | 7 | 1438 (1346-1486; 1326, 1526) | 1607 (1553-1630; 1374, 1724) | 1655 (1627-1687; 1382, 1731) |
|  |  | 2023/24 | SV | 22 | 1210 (1148-1338; 933, 1684) | 1377 (1325-1480; 1082, 1850) | 1395 (1355-1506; 1103, 1857) |
|  |  | **All** | **All** | **54** | **1436 (1331-1513; 923, 1881)** | **1594 (1487-1647; 936, 2020)** | **1621 (1517-1676; 938, 2102)** |
|  | Post-brood<sup>6</sup> | 2019/20 | SV | 2 | 1083 ( 854-1311; 625, 1540) | 1297 (1106-1487; 915, 1678) | 1353 (1170-1537; 986, 1720) |
|  |  | 2020/21 | SV | 4 | 1163 ( 848-1469; 738, 1550) | 1466 (1203-1719; 1049, 1842) | 1564 (1369-1788; 1302, 1940) |
|  |  | 2021/22 | SV | 3 | 583 ( 576- 876; 569, 1170) | 1294 (1108-1376; 922, 1458) | 1471 (1443-1512; 1415, 1553) |
|  |  | 2021/22 | UT | 3 | 1588 (1437-1709; 1285, 1830) | 1953 (1837-2084; 1720, 2215) | 1989 (1890-2131; 1791, 2272) |
|  |  | 2022/23 | SV | 19 | 674 ( 537-1468; 482, 2325) | 941 ( 803-1654; 612, 2483) | 1000 ( 841-1733; 628, 2644) |
|  |  | **All** | **All** | **31** | **1037 ( 742-1440; 482, 2325)** | **1323 (1112-1689; 612, 2483)** | **1397 (1218-1749; 628, 2644)** |
| GPS | Incubation | **2022/23** | **SV** | **27** | **800 ( 630-1180; 388, 1626)** | **1119 ( 794-1356; 558, 1746)** | **1153 ( 804-1450; 597, 1782)** |
|  | Brood-guard | **2022/23** | **SV** | **8** | **540 ( 474- 552; 314, 726)** | **601 ( 563- 652; 345, 787)** | **608 ( 571- 656; 381, 792)** |
|  | Post-brood<sup>7</sup> | 2022/23 | SV | 11 | 595 ( 494- 674; 474, 895) | 784 ( 686- 969; 511, 1418) | 801 ( 694-1010; 513, 1447) |
|  |  | 2023/24 | JT | 24 | 639 ( 592- 725; 529, 1314) | 1065 ( 754-1338; 645, 1758) | 1075 ( 783-1364; 653, 1781) |
|  |  | **All** | **All** | **35** | **617 ( 543- 700; 474, 1314)** | **925 ( 720-1153; 511, 1758)** | **938 ( 738-1187; 513, 1781)** |

<sup>1</sup> Shortest distance from the colony, given movement constraints (see Methods). <sup>2</sup> For GLS- and GPS-tracked birds, distances refer respectively to all at-sea locations or foraging locations only. <sup>3</sup> JT = Jutulsessen, SV = Svarthamaren, UT = Utsteinen. <sup>4</sup> Median, 95<sup>th</sup> percentile and maximum calculated at the individual level, then aggregated across seasons/stages/colonies. <sup>5</sup> May be inflated due to geolocation error. <sup>6</sup> Observed Feb 5<sup>th</sup> to Mar 16<sup>th</sup>. <sup>7</sup> Observed 19<sup>th</sup> Jan -1<sup>st</sup> Mar.

## 3. Results

### 3.1 Sample sizes and breeding phenology

We deployed GLS loggers on 53 birds, the majority (77 %) at Svarthamaren, and retrieved data from 34 (Table 1), tracked for a median of 307 d (IQR 301 - 351 d; range 287 - 1462 d). Data coverage was biased towards the pre-laying exodus, and the last two study seasons. We deployed GPS loggers on 75 birds (Table 1) and retrieved data 60 birds, the majority from incubation at Svarthamaren in 2022/23 and post-brood at Jutulsessen in 2023/24 (Fig. 2). Birds were GPS-tracked a median of 2 trips (IQR 1-3; max 9) over 10.8 d (IQR 7.2 – 19.7; range 2.9, 40.3) at an interval of 30 min. (IQR 30 – 60; range 30, 120). Across colonies, females were 16 g (95% CI 4-28 g) lighter than males (Tables S2 & S3) and female mass was lower at Jutulsessen (232 ± 24 g) than at Svarthamaren and Utsteinen (∼ 258 ± 32 g at both).

**Fig. 2.**
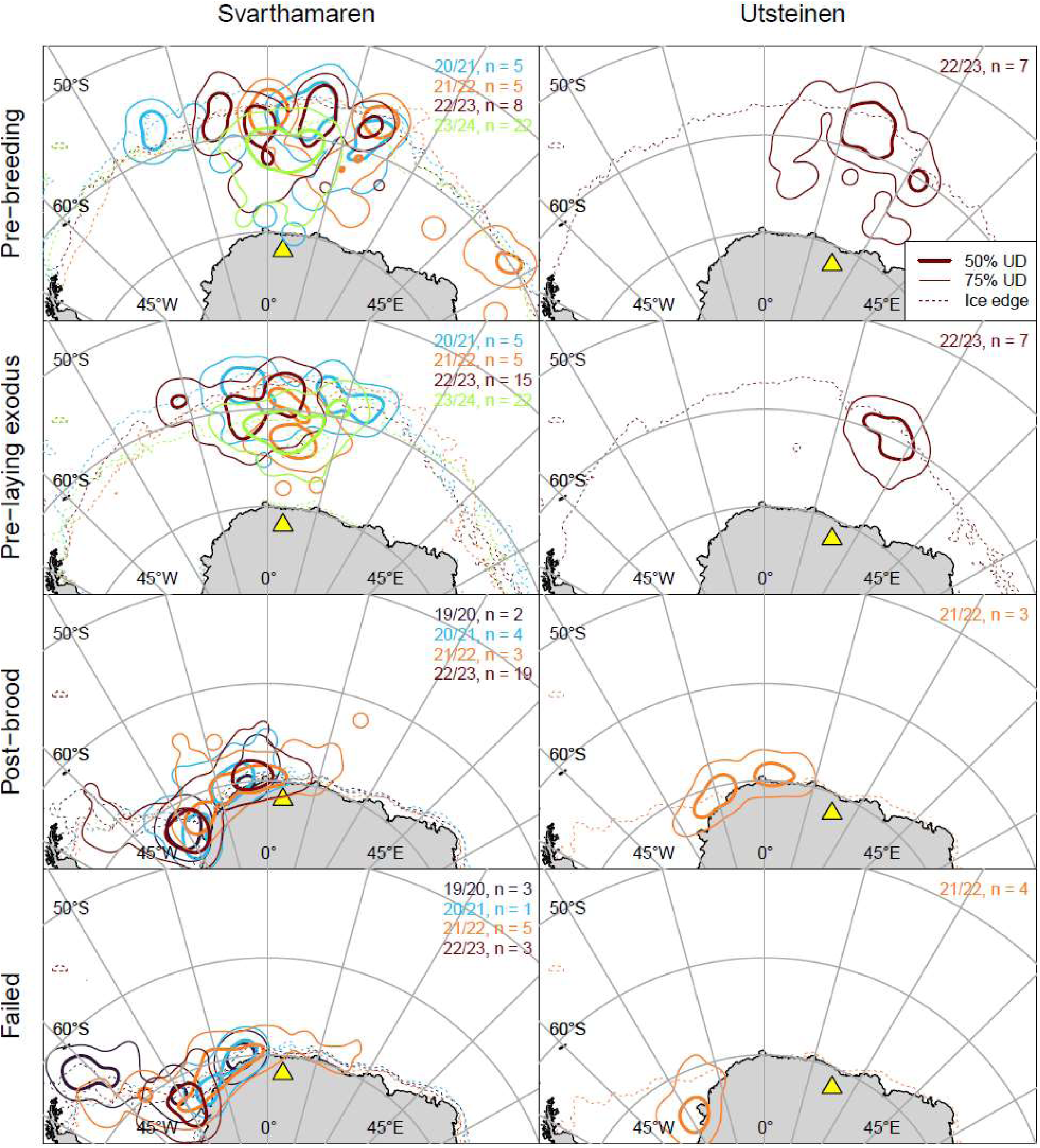
Home ranges of snow petrels tracked with GLS during three breeding stages (rows) from two colonies (columns) in Dronning Maud Land in five breeding seasons (colours). N = number of birds tracked in each stage/season, plus failed breeders. Yellow triangles show study colonies.

Based on GLS data, the pattern of first nest attendance was similar across individuals and colonies: excluding one individual that attended its nest from 14-16 October, the first full day spent on the nest was 07 November ± 3 d (range 03 November – 16 November) and did not differ between Svarthamaren and Untsteinen (*t*(32) = 0.578, p = 0.567). Most individuals experienced 24 h daylight for 1 d prior to this, suggesting that they arrived at the latitude of the colony a day before entering the nest cavity. The majority (82%) of birds spent one bout (9.1 ± 2.8 d) at the breeding site prior to incubation, with males spending 3.8 d longer than females (*t*(18) = -7.217, p < 0.001). Respectively, 12 and 6 % of birds undertook a second and third pre-laying bout (2.5 ± 0.8 d and 3.5 ± 2.1 d, respectively). The first bout on the nest was 3.8 d longer among males than females (*t*(18) = -7.217, p < 0.001). The pre-laying exodus began on 18 November ± 5 d (range 11 November – 04 December) and the first incubation shift on 04 December ± 4 d (range 30 November 30 – 11 December 11). Direct nest monitoring at Svarthamaren (n = 78 nests) recorded laying on 04 December ± 4 d and hatching on 14 January ± 4 d. Mean (range) trip durations of GPS-tracked birds were: incubation 7.0 ± 2.5 (2.9-11), brood-guard 3.8 ± 0.7 (2.9-4.7) and post-brood 4.6 ± 1.2 (2.1-7.5) d.

### 3.2 Space use and foraging range

Most GLS-tracked snow petrels spent the pre-breeding period and pre-laying exodus in a 45° sector north of the respective colonies (Fig. 2). Core home ranges in both stages were centred around 100 km south of the ice edge. Within these stages, core home ranges of birds from Svarthamaren overlapped partially among seasons (*HROI* ranges from 0.04 to 0.39 for pre-breeding and 0.02 to 0.46 for pre-playing exodus, Fig. S3). During spring 2022, when birds from both Svarthamaren and Utsteinen were tracked, core home ranges of the respective populations overlapped slightly during pre-breeding (*HROI* = 0.12) but not during the pre-laying exodus (Fig. 2). Within these stages, median foraging range *d*_*u*,50_ (Table 2) did not differ significantly between colonies (Svarthamaren vs Utsteinen pre-breeding, *t*(13) = 0.234, p = 0.819; pre-laying exodus, *t*(19.3) = 0.135, p = 0.894) so we aggregated across colonies for comparisons among years. In addition, as individual repeatability within life-history stages for birds that were tracked in multiple seasons was low (*R* for *d*_*u*,50_ ≤ 0.3), we treated repeated measures on individuals across years as independent. Within these stages, *d*_*u*,50_ varied significantly among years (Table S4). Prior to breeding, birds used locations approximately 540 km closer to their colonies in 2023/24 than in other seasons. During the pre-laying exodus, *d*_*u*,50_ was approximately 240-310 km lower after 2020/21. For birds tracked during both pre-breeding and the pre-laying exodus, *d*_*u*,50_ was ∼170 km lower (*t*(14) = 3.361, p = 0.005) during the former stage in 2022/23 but did not differ between these stages in 2023/24 (*t*(21) = -0.459, p = 0.651). Among 20 known-sex birds, males foraged closer to the colony than females by ∼220 km during the pre-laying exodus (*t*(18) = 2.439, p = 0.025) and ∼210 km during the pre-breeding period, but the significance of the latter was marginal (*t*(18) = 2.085, p = 0.052).

Birds GLS-tracked from Svarthamaren during late post-brood (05 February to 16 March) concentrated in Weddell Sea coastal waters, from approximately 0 to 38°W (Fig. 2). Home range overlap was relatively high among years (*HROI* range 0.29 – 0.72, Fig. S3) but sample sizes were insufficient to test for differences in *d*_*u*,50_ among colonies and seasons (Table 1). Six birds tracked from Utsteinen and Svarthamaren in late post-brood in 2022 used a similar area (*HROI* = 0.66). Failed breeders used areas further west, centred either in the southeast Weddell Sea or in 2020, the northwest Weddell Sea (Fig. 2).

Paths followed by GPS-tracked birds during overland commuting segments of foraging trips were complex (Fig. S4) but headings on outward and inward overland commutes were approximately northwest and south, respectively (Table 3), crossing the coast on average 4.1-4.8° further west on the outbound than return commute (paired *t*-test (one trip per bird) Jutulsessen *t*(23) = -6.163, p = <0.001; Svarthamaren *t*(32) = -6.239, p = <0.001). Having reached the coast, they largely avoided crossing land or ice shelves again until their inward commute, excepting a few instances when they crossed the Troll Ice Tongue west to east.

**Table 3.** Headings of GPS-tracked snow petrels on overland legs of foraging trips and the longitudes of the locations *s*_l_ and *s*_2_ at which they crossed the coast on outward and inward commutes from three colonies in Dronning Maud Land.

| Colony<br>(longitude) | Leg (n birds) | Mean heading<br>($\rho$)<sup>1</sup> | Median (IQR; range) coast<br>crossing longitude |
| --- | --- | --- | --- |
| Jutulsessen<br>(2.5°E) | Outward (24) | 313.1° (0.97) | 2.9°W (4.0 – 2.0°W; 9.4°W, 2°E) |
|  | Inward (24) | 166.5° (0.91) | 1.2°E (2.0°W – 3.7°E; 7.5°W, 5.7°E) |
| Svarthamaren<br>(5.2°E) | Outward (35) | 321.8° (0.92) | 0.5°E (1.9°W – 3.1°E; 6.3°W, 5.6°E) |
|  | Inward (35) | 179.6° (0.94) | 5.3°E (4.4°E – 6.3°E; 1.7°W, 15.6°E) |
| Utsteinen<br>(23.3°E) | Outward (14) | 343.5° (0.97) | 21.3°E (20.4 – 22.6°E; 18.5°E, 24°E) |
|  | Inward (13) | 199.6° (0.99) | 25.5°E (24.6 – 27°E; 22.8°E, 27.5°E) |

<sup>1</sup> $\rho$ = mean resultant length

Where sample sizes were sufficient for testing (Svarthamaren, incubation, 2022/23; Jutulsessen, post-brood, 2023/24), space use and foraging range based on GPS data did not differ significantly between the sexes (Table 4, Fig. S5). Hence, we pooled sexes in subsequent analyses. Among birds GPS-tracked in 2023/24 (early December to late February), 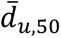 was greatest during incubation, followed by post-brood and brood-guard (Table 2). Differences between these stages were all significant except brood-guard vs. post-brood (Table 5). During incubation, birds foraged mainly <200 km from the coastline, between 7°E and 15°W but also made long trips over a wide area as far north as 58° S, up to 1782 km from the colony (Fig. 3). During brood-guard, they foraged in a similar core area (*HROI*_IN,BG_ = 0.62, p = 0.506) but concentrated nearer the coast, between 0 and 8° E, so UDs differed significantly overall (*BA*_IN,BG_ = 0.62, p = 0.012). During post-brood, they used a similar core area (*HROI*_BG,PB_ = 0.64, p = 0.574; *BA*_BG,PB_ = 0.81, p = 0.232) but also made longer trips to the northwest and west, into the Weddell Sea embayment, as far as 28° W, ≤1447 km from the colony. Post-brood birds GPS-tracked from Jutulsessen in 2024 travelled up to 1781 km from their colony and although they used some of the same locations as 2023 post-brood birds from Svarthamaren, core areas of these two groups differed significantly (*HROI*_sv,JT_ = 0.49, p = 0.002; *BA*_sv,JT_ = 0.74, p = 0.001).

**Fig. 3.**
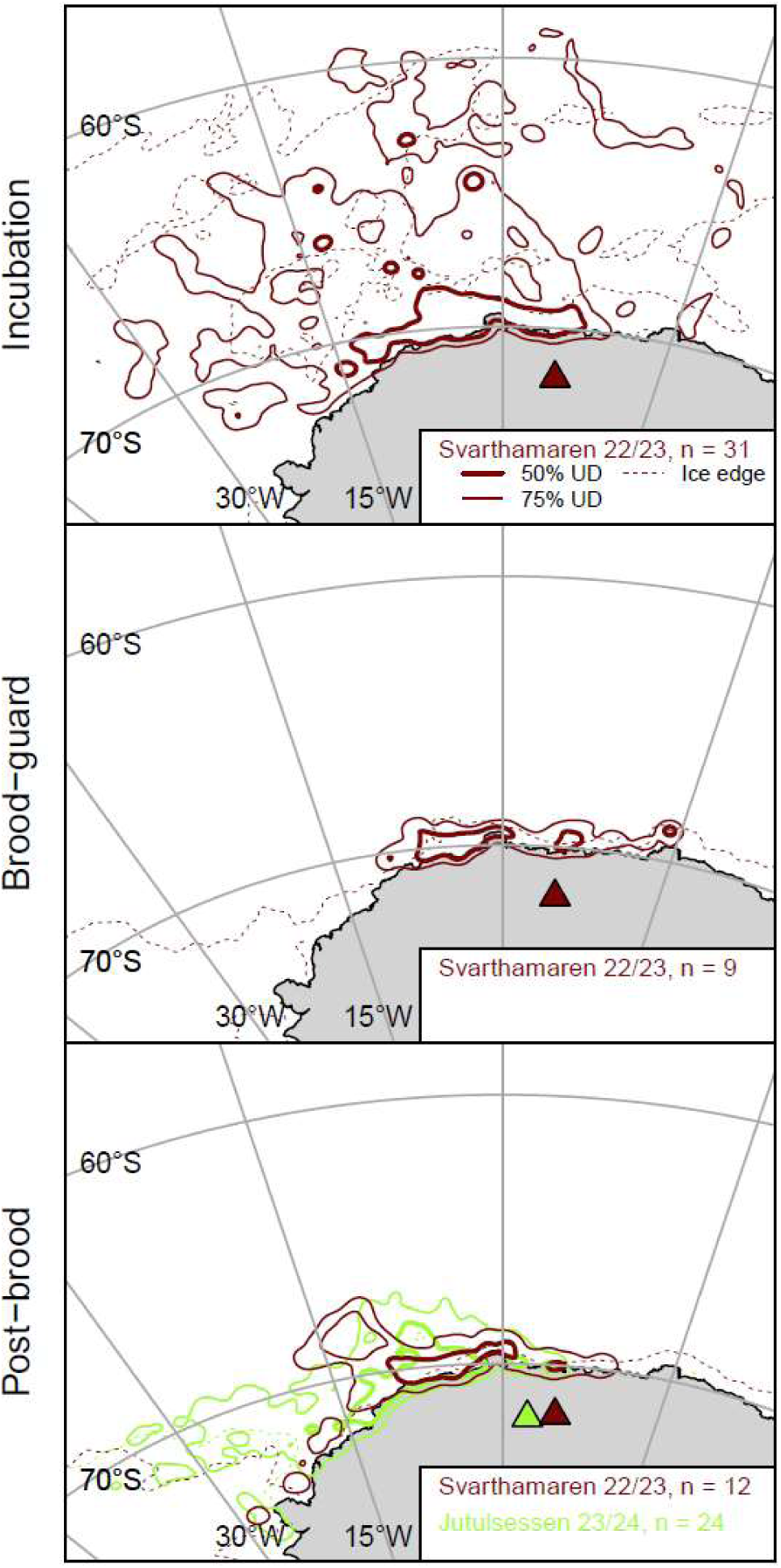
Home ranges of snow petrels tracked with GPS during three breeding stages (rows) from two colonies in Dronning Maud Land in two breeding seasons (colours). N = number of birds tracked in each stage/season/colony and triangles show study colonies.

**Table 4.**
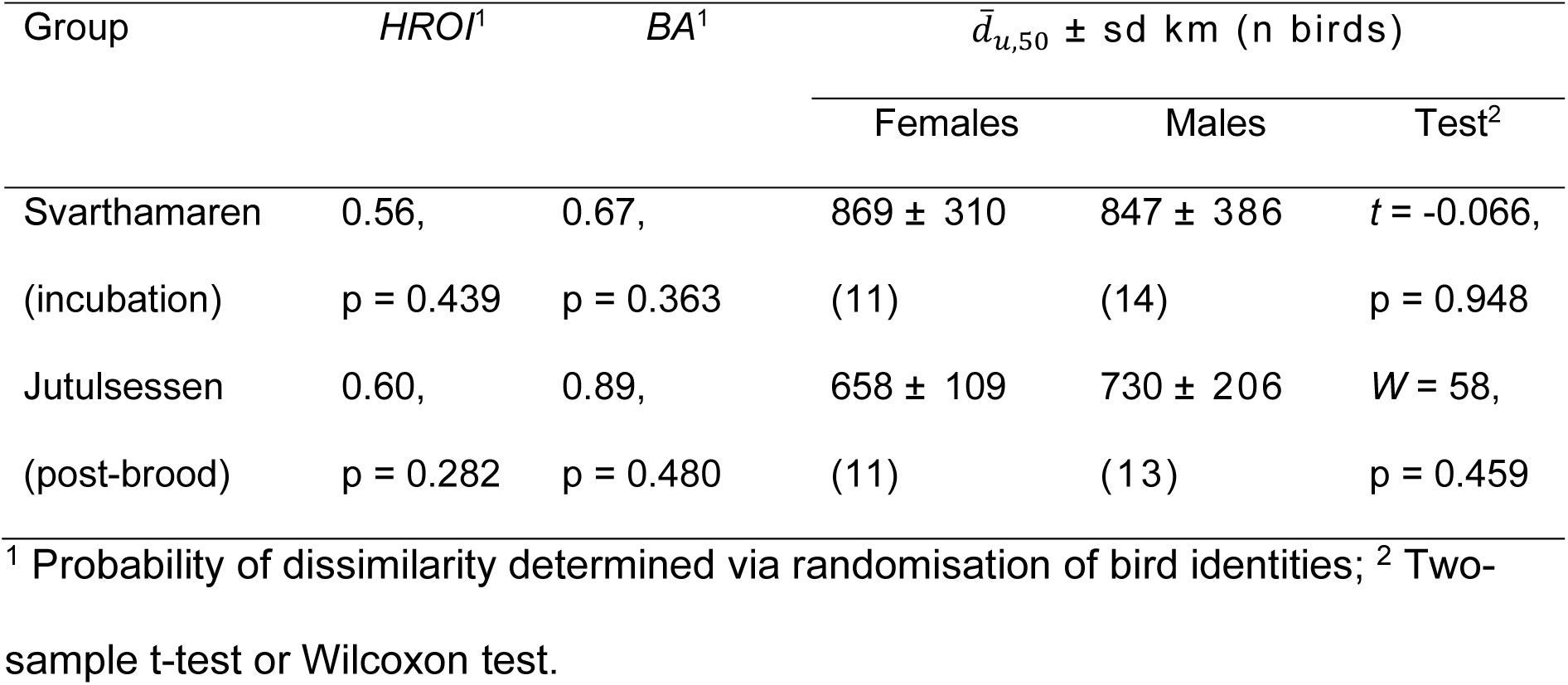
Comparison between foraging distributions and median foraging ranges 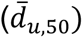 of male and female snow petrels tracked with GPS from two colonies/breeding stages in Dronning Maud Land.

**Table 5.** Fixed effects in a generalized linear mixed-effects model of median foraging ranges *d*_*f*50_ of snow petrels GPS-tracked from Svarthamaren, Dronning Maud Land in 2022/23 as a function of breeding stage.

| Stage | Parameter | SE | $t$ | $P$ | Tukey contrasts $p$ | |
| --- | --- | --- | --- | --- | --- | --- |
|  |  |  |  |  | BG | PB |
| Incubation (intercept) | 0.207 | 0.034 | 6.046 | <0.001 | <0.001 | <0.001 |
| Brood-guard (BG) | 0.237 | 0.051 | 4.627 | <0.001 |  | 0.063 |
| Post-brood (PB) | 0.120 | 0.037 | 3.281 | 0.001 |  |  |

### 3.3 Foraging latitude vs. ice edge latitude

Across breeding stages and seasons, median foraging latitude was strongly associated with median ice edge latitude (Fig. 4), with a linear model fitted to the GLS data showing that birds foraged approximately 2° S of the ice edge. This was also evident during the pre-breeding and pre-laying exodus stages, but not within post-brood. Median foraging latitude estimated from GPS data followed the same trend during incubation, but during chick-rearing it coincided with the latitude of the median ice edge.

**Fig. 4.**
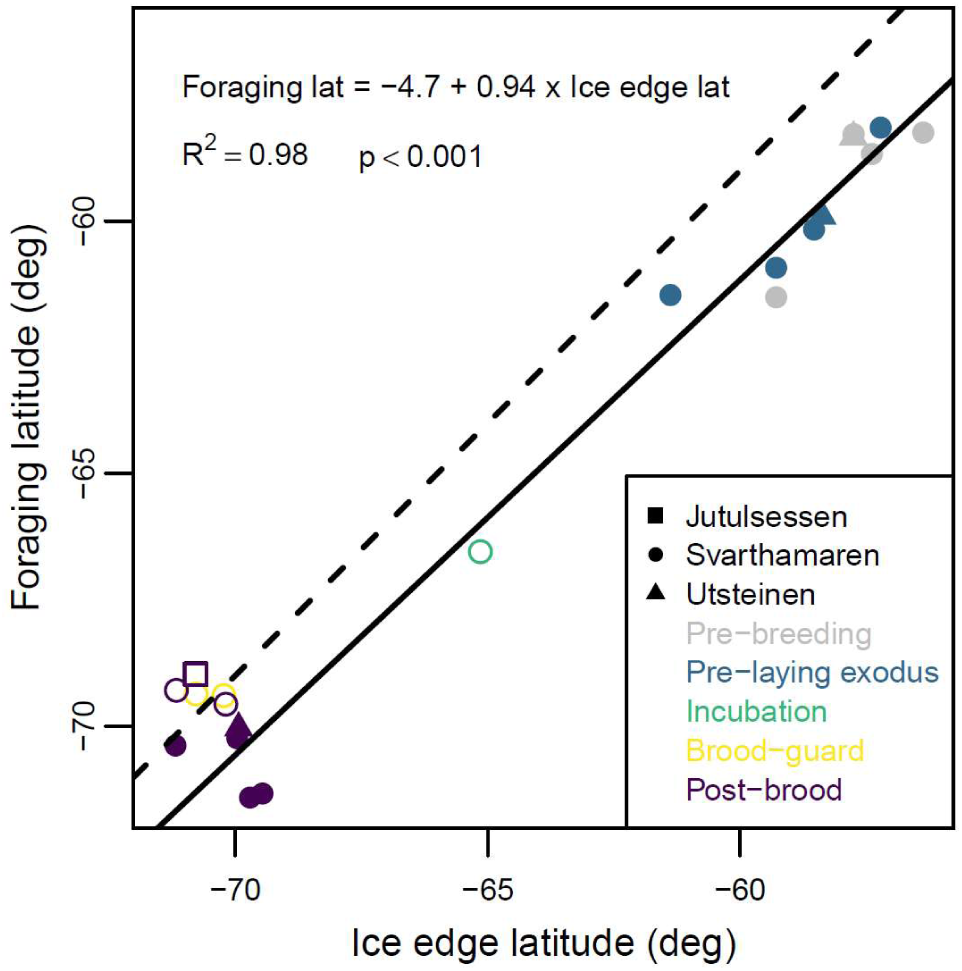
Median foraging latitudes of snow petrels tracked from three colonies in Dronning Maud Land averaged across individuals within stage/colony/season vs. median latitude of the ice edge between 35°W and 25°E during the same periods. The solid line is a linear model fitted to the data from birds tracked with GLS (solid symbols). For comparison, open symbols show data derived from GPS tracking and the dashed line a 1:1 relationship.

## 4. Discussion

### 4.1 Comparison with previous studies and study limitation

Based the distance of breeding sites from suitable foraging habitat, the maximum foraging range of snow petrels has previously been assumed to be at least 480 km [38]. Our data show that even during brood-guard, when birds are most constrained, realised foraging ranges can be ∼25% greater. Indeed, the 95^th^ percentile foraging range of snow petrels, varies from ∼600 km during brood-guard to ∼1600 km during the pre-laying exodus, reflecting the relative central-place constraint associated with different breeding stages and seasonal fluctuations in the location of the MIZ, where snow petrels typically forage. The mean foraging range of snow petrels tracked with GLS from Ile des Pétrels, Terre Adélie during the pre-laying exodus and late post-brood was 2648 ± 1054 (max 4978) km [40]. This is around 1000 km greater than we observed in either stage, but the Terre Adélie dataset may have included failed breeders. In contrast, birds tracked via GPS from the same colony during incubation had a median foraging range of only ∼120 km [figure 1 in 11], ∼600 km less than we observed. In part, this may be due to Svarthamaren being ∼175 km further inland but given the relationship we found between meridional foraging range and sea-ice extent, another likely cause is regional differences in sea ice distribution and dynamics. In December, the median ice edge is approximately 580 km from Svarthamaren but only 40 km from Ile des Pétrels (Fig.4). In addition, intraspecific competition may force birds from DML to forage further from their colonies because regional population size is apparently relatively high compared to Terre Adélie [7].

While it has sometimes been assumed that the breeding distribution of snow petrels is limited by distance to the coast [18, 39], it is more likely that distance to the MIZ is limiting, because this is their preferred foraging habitat [10] (section 4.4). During early chick-rearing, when snow petrels are most constrained and sea-ice extent is near its seasonal minimum, the maximum distance between any currently occupied colony and the median 50% SIC contour is 694 km [77]. This is similar to the maximum foraging range that that we observed for this stage (median 608, IQR 570-655, overall maximum 791 km), so we tentatively conclude that the upper limit of sustainable foraging range during chick-rearing is around 700 km.

However, our results have several potential limitations. Firstly, we used GLS to track birds during the pre-laying and late post-brood stages, and GPS during incubation through to mid post-brood. While the accuracy of GLS tracking is an order of magnitude lower than GPS-tracking [57, 58], GLS errors are approximately bivariate normally distributed [59] so should not bias our estimates of the median foraging range. However, our GLS-derived 95^th^ percentile and maximum forage ranges will be sensitive to outliers and should be treated with caution. For example, during late post-brood, our dataset may have included some failed breeders, which no longer central-place constrained. Secondly, light-based geolocation cannot resolve latitude around the equinoxes and neither latitude nor longitude can be resolved in spatiotemporal regions of 24 h daylight [78][refs]. Pre-breeding was largely unaffected by the equinox (Fig. S1), but the pre-laying exodus occurred when 24 h daylight extended to 64°S, so that locations of any birds foraging in coastal waters would have been unresolved. In practice, we suspect that this would not have caused a large bias because the general pattern (confirmed by GPS tracking) through to the end of incubation was for birds to forage 2° south of the outer ice edge, which in the pre-laying exodus was located between ∼58-61°S (Fig. 2). During late post-brood, 24h daylight could also have resulted in failure to resolve foraging locations far from colonies in the southern Weddell Sea (cf. Fig. S1 and Fig. 2). However, we estimated a greater late post-brood foraging range via GLS compared to early post-brood estimates via GPS, consistent with foraging range increasing over the chick rearing period observed in other species [e.g., 79, 80; see Section 4.2]. Due to these ambiguities in GLS data, we consider the foraging range estimates based on GPS data to be more reliable for post-brood. A final caveat (applicable to most tracking studies) is that although our devices were relatively small [81], the extra mass and drag could have affected foraging range [82]; however, this was more likely to reduce, rather than increase the range [83, 84].

### 4.2 Intrinsic and extrinsic effects on foraging range

Seasonal variation in snow petrel foraging ranges conformed to the pattern typically observed among pelagic seabirds, ultimately reflecting intrinsic breeding constraints [1, 3, 27, 79]. Range was greatest prior to breeding, when birds are least constrained, and during the pre-laying exodus, when they return to sea for approximately two weeks. Lowest ranges were recorded during brood-guard, when parents alternate between attending and feeding the newly-hatched chick, in our study typically swapping every four days.

Flight speed, fasting endurance of adults and chicks, and many other traits scale allometrically in seabirds [85]. Among breeding snow petrels, trip duration is negatively associated with body size [51]. It might therefore be expected that foraging range increases with body size [3, 86]. We studied snow petrels of the ‘lesser’ form [8, 50, 87, 88], and so our results may not be generalisable to the larger form breeding elsewhere, particularly if there is size-mediated competitive exclusion [51].

Size-mediated competitive asymmetry, as well as other sexual differences, can also result in differences between male and female foraging areas and ranges [28, 29, 89–91]. Males in our study were 6-7% heavier than females, which is typical for the species [87, 92, 93]. Our sample of known-sex birds during pre-breeding and the pre-laying exodus was small and the effect of season potentially confounding (Table 1). Nonetheless, our results are consistent with a sex difference in breeding duties suggested by previous, colony-based studies [51, 94]. Among birds tracked in 2022/23 and 2023/24, males foraged around 200 km closer to the colony than females during both pre-breeding and the pre-laying exodus. We assume that this is due to an imperative for males to return earlier to the colony and invest more time in nest defence (also evinced by the greater amount of time they spent at the nest site prior to breeding) and for females to invest more time in egg production [89, 95]. In contrast, and in common with snow petrels tracked from Terre Adélie during incubation [11], we found no evidence of spatial segregation or differences in foraging range between sexes during incubation or chick rearing. This does not preclude niche partitioning between the sexes along axes of habitat (e.g., sea-ice concentration) or diet, as in Barbraud et al. (2021). Habitat partitioning could occur without broad differences in space use or foraging range as sea-ice concentration in the spring and summer in our study area is highly patchy and dynamic due to temperature, winds and currents [42, 96].

Evidence is growing that flying seabirds track particular sea-ice habitat in time and space [35, 97], so sea-ice dynamics might be expected to affect foraging range. We found a strong positive association between the meridional location of the ice edge and foraging latitude, which was around 2° (∼220 km) south of the ice edge - i.e., within the MIZ. This relationship was evident not only within but also across breeding seasons (Fig. 4). For example, just prior to colony return and during the pre-laying exodus in 2023, when sea-ice extent was at a record low [98], birds foraged closer to the colony than in previous years (Fig. 2) and in 2021/22 and 2023/24, core home ranges during the pre-laying exodus encompassed the Maud Rise, where seasonal sea-ice thinning occurs earlier than in the surrounding pack (Fig. 1). Other GLS studies suggest that snow petrels track the ice edge over winter (c.f. Delord et al. [40], Viola et al. [99], Fetterer et al., [60]), and at-sea observations indicate that they do so at both large and fine spatiotemporal scales [36, 37, 100]. In our study, the relationship between the latitudes of snow petrel and the ice edge broke down during chick-rearing (mid-January, Fig. 4), when open water occurs all the way to the coast (Fig. 1). From this time, birds increasingly moved west following the MIZ, which continues to retreat in the eastern Weddell Sea, until the summer minimum occurs in February (cf. Figs. 1, 2 and 3).

Given that sea ice in the vicinity of most snow petrel colonies around Antarctica retreats broadly towards colonies, we hypothesise that breeding schedules and therefore potential foraging ranges, are synchronised to sea-ice dynamics, such that chicks hatch when suitable foraging habitat is closest to the colony [101], then during the chick-rearing period, adults take advantage of the peak in secondary production that typically follows sea-ice breakup by several weeks to months in our study area [35, 97]. Matching of breeding schedules to maximise access to resources during the most energetically demanding phase may be particularly important for snow petrels because they breed at such high latitudes that their breeding season is relatively short [102]. Further investigation of the resource-tracking hypothesis should explore the effects of seasonal variation in habitat availability on habitat selection [97]. In the meantime, we caution that our study was undertaken during a four-year period in which summer sea-ice extent was anomalously low and the rate and timing of summer melt unusually rapid and early [45, 103, 104]. The foraging ranges we observed may have been lower than in periods of more extensive sea ice.

Among central-place foraging seabirds, both foraging range and distribution can be affected by density-dependent competition [31, 32]. Within stages, we did not observe large differences in foraging range among colonies, possibly because numbers of birds breeding at or near those sites were of a similar magnitude [1000s; 7, 77]. However, where we were able to compare core home ranges among colonies, these were either partially or fully segregated prior to incubation. This is not necessarily due to competition: segregation could have arisen from birds commuting between their colonies to the nearest available habitat patch, which during spring and early summer is usually immediately to the north. By post-brood, birds from all three study sites had partially overlapping core home ranges, presumably because large patches of intermediate SIC occurred only near the coast, west from the prime meridian into the Weddell Sea embayment (Fig. 1). This is consistent with studies showing that sharing of space tends to occur at similar distances to colonies and when food availability is superabundant [105]. Indeed, density-dependent competitive effects in our study system may be less than at lower latitudes because the productivity of Antarctic waters during the summer is so high [106]. Despite this, there was significant dissimilarity between the post-brood core areas of birds from Svarthamaren and Jutulsessen. Given that these colonies are only 100 km apart, this may have been due to competition or simply to the two groups being tracked in different years.

The other extrinsic effects evident in our data were those imposed by wind and topography on movement. It was beyond our scope to investigate these in detail, but we attempted to account for them in analysis via several assumptions. Firstly, that once at sea, commuting snow petrels avoided crossing land until they were returning to the colony. This assumption is orthodox [41] and well supported by our data. Route choice during the relatively long commute between the colony and the coast was more complex. Typically, outward commutes over the ice sheet were more westerly than the reciprocal headings of inward commutes. Other Antarctic fulmarine petrels also behave in this way, presumably to optimise movement through the strong prevailing easterly winds of the Antarctic coastal zone [72, 107]. In short, it hypothesised that this strategy allows wind drift on the way out but compensates for it on the way back. In order to estimate realised foraging ranges taking into account these movement constraints, we approximated distance on overland commutes by beelines to coast crossing locations observed via GPS-tracking and assumed that these were similar for birds tracked with via GLS. Although these assumptions are expedient, further observation and analysis of the effects of wind on travel costs and route optimisation strategies [27, 108] may result in refined foraging range estimates and projections. Despite these reservations, we are confident that distances as defined in our study are a more realistic indication of the actual commuting distances than the beeline distances used in many previous studies. For example, beeline and biological distances between the colony and the location illustrated in Fig. S2 are 734 and 1065 km, respectively, illustrating that the former can greatly underestimate potential foraging range.

### 4.3 Implications for interpretation of paleo-climate and occupation histories from stomach-oil deposits

Interpretation of palaeoclimatological and ecological proxies from stomach-oil deposits involves assumptions about where the snow petrels could have foraged [5, 6, 43]. Previously, these were based on sparse data that typically underestimated potential foraging ranges. Using our results, we have projected potential foraging ranges for all breeding sites in our study region (see Supplementary Materials), which we hope will aid interpretation of proxies derived from stomach oil deposits. These show that areas accessible to snow petrels, and therefore over which stomach oil deposits potentially integrate sea ice and other environmental conditions could be as large as 5 million km^2^ (Fig. 5a). However, our results also show that foraging ranges contract by a factor of two to three over the breeding cycle. Hence, it is important to consider how stomach oil deposition might vary seasonally. If adults and chicks spit oil at a similar rate throughout the breeding season, deposits would integrate paleo-environmental information over a wide range of distances and sea-ice conditions, but with a bias towards locations <900 km from the colony (Fig. 5b). Alternatively, oil deposition could occur in one or more short pulses, the most likely being between first return to the colony and early incubation, when competition to establish (or reestablish) and defend nest sites and mates is intense [8, 94]. In this scenario, accumulated stomach oil would reflect environmental conditions prior to laying, when potential foraging ranges exceed 1400 km and sea-ice extent is near its seasonal maximum [109]. Moreover, accumulated oil could originate predominantly from males, which are more active in nest acquisition and defence and take the first incubation shift [94, 110]. This is relevant because, as discussed above, males forage closer to the colony than females during pre-breeding and the pre-laying exodus. Deposition could also vary due to predation pressure from skuas, which in Terre Adélie peaks in early post-brood [14].

**Fig. 5.**
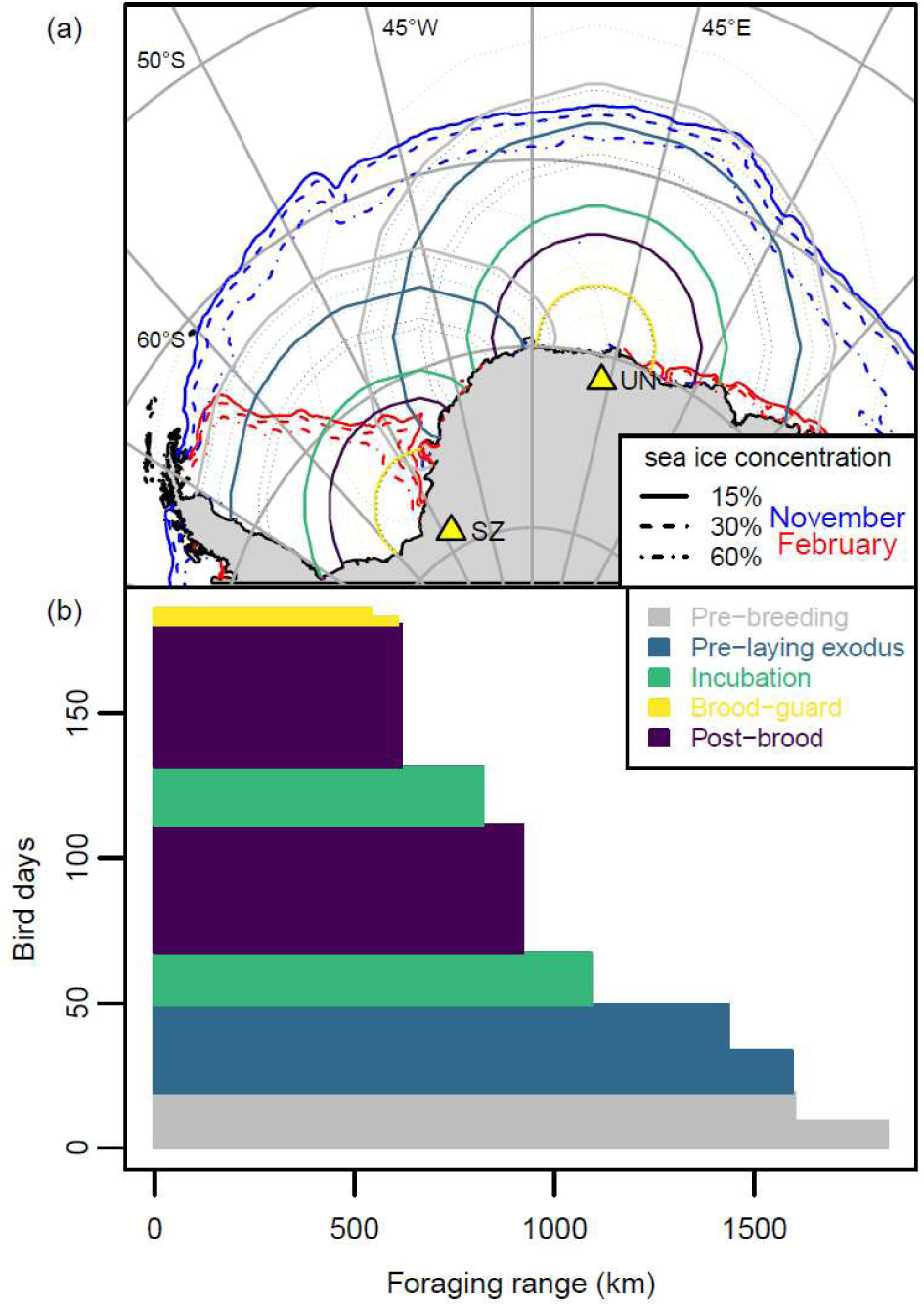
(a) Potential 95^th^ percentile foraging ranges (with 95% confidence intervals) of snow petrels from Untersee Oasis (UN) and Station Z.508 (SZ) during all breeding stages, plus 30 y median sea ice concentration contours. (b) Number of bird days that areas within different distances from the colony are accessible to snow petrels over the breeding season based on this study (bird days = length of breeding stage x number of parents that forage simultaneously during that stage (i.e. one or two) x 0.5 or 0.45 for the median and 95^th^ percentile ranges, respectively.

On balance, we suggest deposition probably peaks early in the breeding season, during courtship and subsequent defence of nests from non-breeders, declining thereafter. Deposits therefore probably predominantly reflect sea-ice conditions during spring and early summer. These assumptions could be tested by directly measuring seasonal variation in deposition rates. The interpretation of dietary signals as indicators of sea ice conditions (e.g. Berg et al., 2019; McClymont et al., 2022) will also need to consider that foraging effort is unlikely to be evenly distributed throughout the potential home range, but is probably patchily distributed due to seasonally changing sea ice conditions, prey availability and possibly competition.

Assuming that snow petrel breeding distribution is limited by both distance to the MIZ and availability of exposed rock for nesting [10], our results can provide new insights into past, present and future nest site occupation patterns [18, 111]. Today, exposed nunataks occur throughout the coastal zone of the Weddell Sea (Fig. 6), but nesting does not apparently occur in eastern Palmer Land or the Pensacola Mountains, where closed, multiyear ice persists in the western Weddell Sea during summer [109]. Snow petrels nest at low densities in the Theron Mountains and possibly the Shackleton Range, where we show that the MIZ of the outer ice edge is beyond the range of birds during the early breeding season (Fig. 6a). Presumably they forage in or at the margins of coastal polynyas [39], such as the Eastern Weddell Polynya, which persistently occurs adjacent to the Brunt Ice Shelf [112, 113]. During chick-rearing, the intermediate sea ice cover occurs over wider but spatiotemporally patchier areas (Fig. 6b), so that accessibility of suitable foraging habitat integrated across the entire breeding season is limiting.

**Fig. 6.**
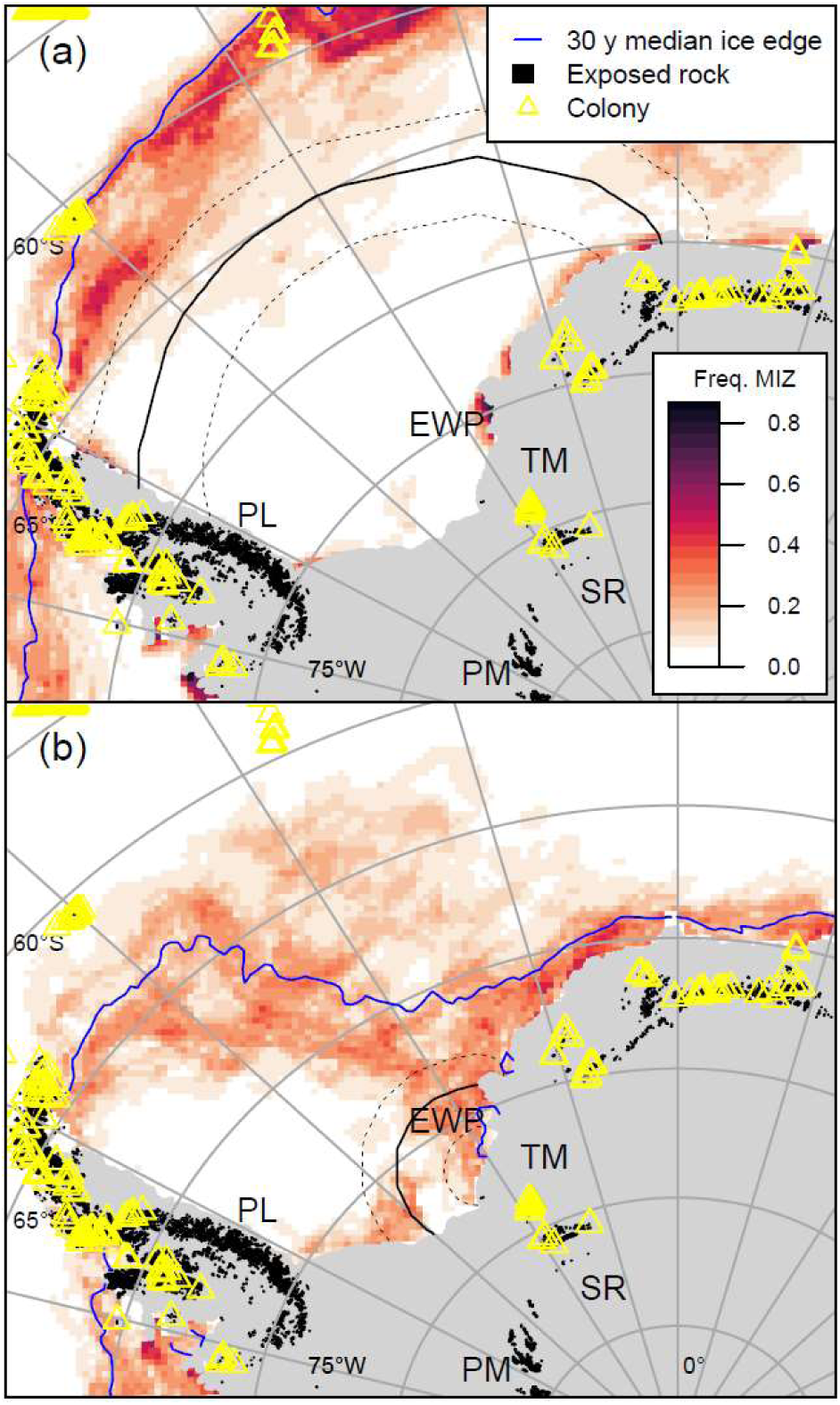
Potential foraging ranges of birds from Station Z.508 (solid yellow triangle), Antarctica during (a) the pre-laying exodus and (b) brood-guard, plus the 30-y mean frequency of occurrence of the Marginal Ice Zone (MIZ - sea-ice concentrations between 30 and 60 %). Station Z.508 is the site furthest from the coast in the Weddell Sea region for which there is robust evidence of breeding [7, 114]. EWP, Eastern Weddell Polynya; PL, Palmer Land; PM, Pensacola Mountains; SR, Shackleton Range; TM, Theron Mountains.

Availability of breeding habitat and access to suitable foraging habitat were thought to be more limited during the last glacial stage [18, 111]. However, snow petrels occupied some colonies in DML throughout this period [16, 17, 44], despite the summer ice edge being >2,000 km north [22, 115, 116]. Our data suggest that this is likely beyond their brood-guard foraging range, supporting the hypothesis that foraging was sustained by polynyas [5, 6, 39]. These could have occurred not only along the coast [117] but also offshore, e.g., over the Maud Rise [118], which is just within the observed brood-guard range of existing colonies in the west of our study area (cf. Fig. 1a and Fig. 6a). Another possibility is that birds foraged in leads, which tend to occur predictably where tidal currents and bathymetry interact, such as at the shelf break [119].

During the transitions between glacial and interglacial stages, which occurred on timescales of tens of thousands of years, there may have been differences in the timing of accessibility to foraging habitat and breeding habitat. For example, retreat of the summer sea ice from its glacial maximum began ∼23 ka in the Atlantic sector of the Southern Ocean [115], but ice-sheet mass loss and warming associated with deglaciation started later, from 20 ka [22, 120]. Under this scenario, foraging habitat may have been available but potential nest sites may still have been ice-covered, especially given that ice sheet thinning occurs later in our study region mainly from ∼10-8 ka [121]. Alternatively, foraging habitat may have become accessible some time after breeding habitat was exposed [122]. This would have introduced latency between breeding habitat exposure and its occupation by snow petrels, which should be borne in mind when inferring ice sheet thinning from stomach oil deposits [18, 44].

Currently, the Southern Ocean is thought to be undergoing a regime shift from a positive to negative trend in summer sea-ice cover [123], exacerbated by anthropogenic climate change [124]. In the future, it is likely the retreat of ice shelves and terrestrial ice sheet margins caused by anthropogenic climate change will impact snow petrels. In particular, foraging habitat (i.e., intermediate sea-ice cover) will become less extensive and seasonally persistent in some areas, whereas breeding habitat (exposed rock) may become more extensive. Hence, snow petrels may be expected to colonise breeding habitat currently at the margins of accessibility to the MIZ and abandon areas where it becomes inaccessible. In our study area, this would result in a southwest shift in breeding distribution, perhaps with areas such as the Pensacola Mountains being colonised for the first time. However, changes in sea-ice phenology [125, 126] and the wider marine ecosystem [34, 127] and ultimately the loss of summer sea ice [21], may have unsustainable impacts on snow petrels.

## 6. Conclusions

Previously, lack of data on the foraging ranges of breeding snow petrels has led to uncertainty in the interpretation of proxies of paleoclimate derived from ancient stomach oil deposits accumulated around snow petrel nests [5, 6, 43] as well as the environmental factors limiting their current and past breeding distribution [25, 26]. Our study shows that not only is foraging range greater than hitherto assumed, but it is also very variable seasonally. Median foraging range was greatest (1436 km) at the start of the breeding cycle during the pre-laying exodus, and least (540 km) during the brood-guard period in the middle of the cycle. A strong correlation between foraging latitude and the latitude of the outer ice edge, plus the known affinity of snow petrels for intermediate sea conditions [9–11], suggests that the decrease in foraging range is due to the seasonal retreat of the pack ice. Indeed, we hypothesise that the breeding schedule of snow petrels is adapted so that brood-guard, when birds are most temporally constrained by chick rearing duties, occurs when the Marginal Ice Zone occurs closest to the colony. Based on our results, and assuming that snow petrel predominantly deposit stomach oil early in the breeding season, when nest defence is most intense, paleoclimate proxies derived from these deposits likely reflect sea ice conditions during spring, when ice cover is near its seasonal maximum. In addition, our results support the hypothesis that breeding at some sites bordering the Weddell Sea may only be feasible due to coastal polynyas or areas of recurrent leads providing foraging habitat south of the outer Marginal Ice Zone [39].

## 7. List of abbreviations

ASPA: Antarctic Specially Protected Area
BA: Bhattacharyya’s Affinity
DML: Dronning Maud Land
GLM: Generalised Linear Model
GLMM: Generalised Linear Mixed-effects Model
GLS: Global Location Sensors
HROI: Home Range Overlap Index
MIZ: Marginal Ice Zone
SIC: Sea Ice Concentration

## 8. Declarations

### Ethics approval

Tracking was approved by the Norwegian Animal Research Authority (permits 21511 and 28555) and Durham University Animal Welfare and Ethical Review Body. It was carried out under permits issued by the UK Foreign and Commonwealth Development Office (13/2021-2022, 08/2022-2023, 04/2023-2024).

### Consent for publication

Not applicable

### Availability of data and materials

The tracking datasets analysed during the current study are available in the BirdLife Seabird Tracking Database https://data.seabirdtracking.org/, dataset ID tbc.

### Authors’ contributions

EW: Conceptualization, Formal analysis, Investigation, Methodology, Roles/Writing - original draft; EM: Conceptualization, Funding acquisition, Writing - review & editing; SD, EH, AR, HR, WG: Investigation, Writing - review & editing; VSB: Formal Analysis, Writing - review & editing; RP: Conceptualization, Data curation, Funding acquisition, Investigation, Writing - review & editing.

## Acknowledgements

We are grateful to the staff at Troll (Norwegian Polar Institute) and Princess Elisabeth (International Polar Foundation) research stations for facilitating data collection.

## 9. Electronic supplementary material

**Supplementary Material 1** – Supplementary Methods, Supplementary Figures S1-S5, Supplementary Tables S1-S4.

**Supplementary Material 2** – Shape files and plots of projected potential foraging ranges of snow petrel colonies in the study area.

## S1. Supplementary Methods

### S1.1. GLS data processing

GPS tracking showed that during incubation birds travelled at least as far north as 59°S (see Results). During midsummer, sea ice recedes to the coast of Dronning Maud Land, above the Antarctic circle making light-based geolocation impractical (Fig. S1). Geolocation is again practicable in late summer, from the date when sunset/sunrise again occurs at latitudes at which snow petrels forage (approx. February 5th), to two weeks before the autumn equinox (March 16th), when geolocation is impractical due to equal daylength across latitudes.

**Fig. S1.**
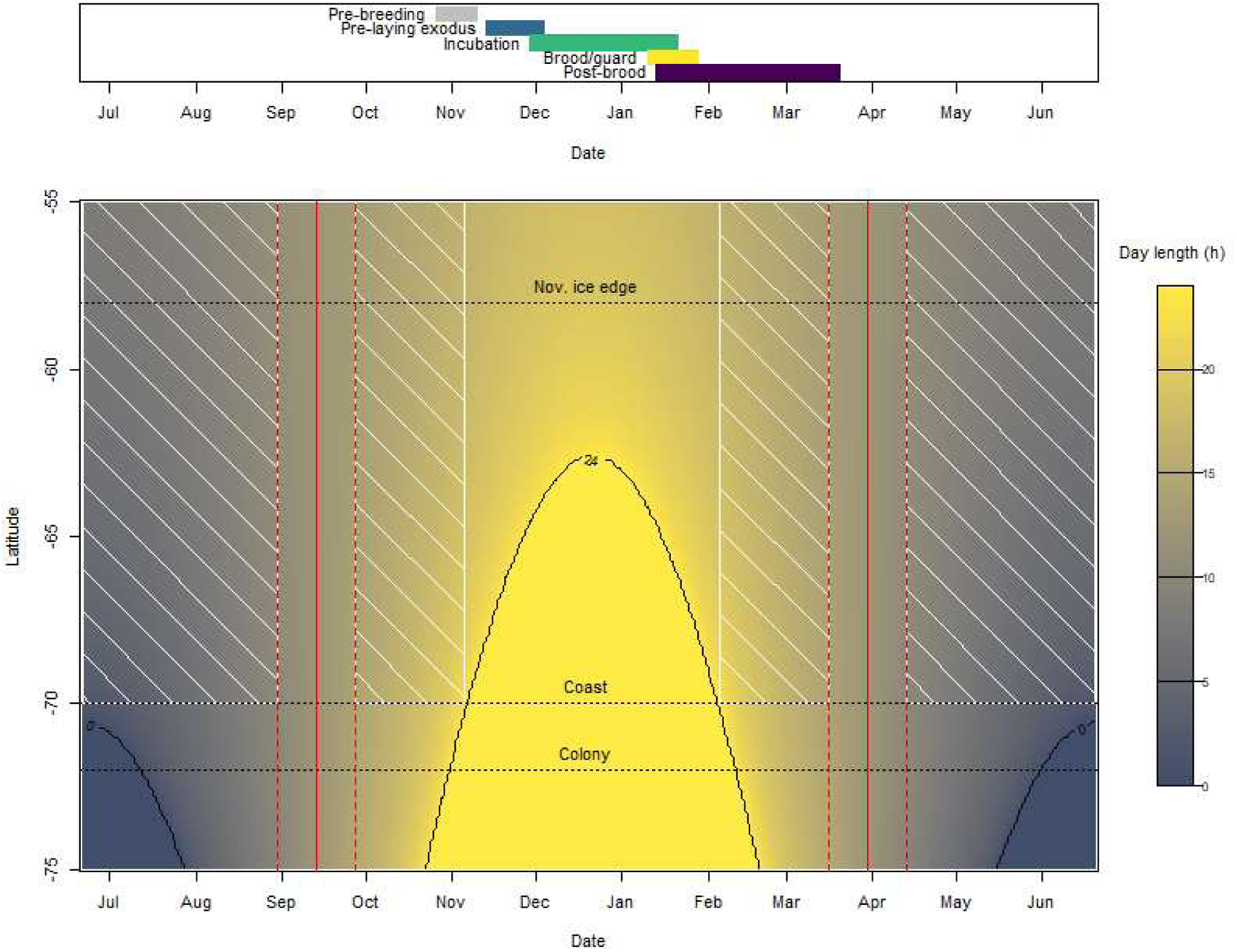
Breeding phenology of snow petrels (top panel) and variation in day length with latitude and time of year and its effect on the tractability of light-based geolocation via the threshold method (bottom panel). Sunrise and sunset are defined here as occurring when the sun is 4.1° below the horizon, the mean values used in our analysis. Red lines show the equinoxes, +/- 14 days; yellow the zone in which 24 h daylight occurs; and the black horizontal dashed lines the approximate latitudes of the November ice edge, the coast of Dronning Maud Land and the study colonies. Areas hatched white are the spatiotemporal regions within which bird locations occurring as far south as the central Dronning Maud Land coast could theoretically be resolved.

Following the protocol developed by Bråthen et al. [1] to analyse geolocation data for mid to high latitude seabirds [2], we used the threshold method to estimate bird locations from the timing of local twilights recorded by the GLS loggers [3, 4]. We identified twilights by assuming that these occurred when light levels crossed a threshold of 11 lux. Following Lisovski and Hahn [5], we fitted polynomial regression models to twilight times, identifying and removing outliers. We assumed that twilight occurred when the sun was at a fixed angle below the horizon but that this could vary between loggers and years [6]. Using methods adapted from Hanssen et al. [7] and van Bemmelen et al. [8], we determined the most appropriate solar angle for each logger/year by visual inspection of plots of latitude vs. time, assuming that the appropriate angle minimised discrepancies in latitudes on either side of the equinoxes. Following this calibration procedure, the mean solar angle was 4.1 ± 0.4° below the horizon. We then estimated locations following Lisovski and Hahn [5] and applied a multi-stage filtering process to reduce errors [1]. Firstly, we removed locations occurring within ±14 days of the equinoxes when day length varies little with latitude.

Due to the solar angle being negative, apparent equinox dates are shifted towards the winter solstice [9], so periods removed were centred on average on October 6 (range October 4 - 8) and March 14 (range March 10 - 18). Next, we applied the distanceFilter function in the GeoLight package [5], removing locations resulting in speeds >40 km/h over 12 h, and then the sdafilter function from the argosfilter package [10], removing unrealistic ‘spikes’ – characterised by steps greater than 480 and 960 km combined with turning angles of less than 15 and 35°, respectively. We then removed locations outside the known range of snow petrels (85 – 49° south) and applied a double smoothing procedure following Hanssen et al. [7].

### S1.2. GPS data processing

We split GPS tracks into foraging trips defining these as excursions beginning and ending ≤1 km from the colony, reaching >50 km from the colony and comprising at least three locations. Due to the power management algorithm used by the GPS loggers, which balances location attempts with battery voltage, and variation in the latter due to shading while birds were on the nest and charging while they were in the open, most locations were recorded at either 30- or 60-minute intervals (see Results). To avoid temporal sampling imbalance biasing our estimates of foraging range, we regularised location estimates in time. To do so, we used the aniMotum package [11] to fit a correlated random walk model to each trip and predict locations at 30 minute intervals. We specified the ‘optim’ optimizer, spike removal distances of 5 and 10 km and bivariate normal GPS location errors of ± 20 m, using defaults for all other settings. We used one-step-ahead prediction residuals to check goodness of fit [12].

### S1.3. Molecular sex determination

We determined sex via standard molecular methods, using primers 2550F and 2718R previously employed for snow petrels [13, 14]. We isolated total genomic DNA from feather tips using the protocol and reagents of the E.Z.N.A. tissue DNA kit (Omega Bio-tek; Norcross, GA, USA), additionally adding DTT (Dithiothreitol) with the proteinase K digested samples overnight in a 55 °C water bath. We eluted DNA twice using 50 µL of low EDTA, TE buffer (Thermo Scientific Chemicals; Waltham, MA, USA). We then amplified DNA using Platinum II Taq Hot-Start DNA polymerase (Invitrogen) for 50 cycles with annealing at 50 °C for 25 seconds and extension at 68 °C for 15 seconds and examined PCR products using 1.2% agarose gels. We ran all PCR amplifications and gels with a known female snow petrel and a negative PCR control (water). We designated males as birds that produced the CHD1Z band only (∼650 bp) and females as those that produced the CHD1W band (∼450 bp) additionally or alone. We reran samples that initially produced no or very faint bands using differing amounts of DNA until sex was unambiguously established.

### S1.4. Approximation of biological distance

We approximated the biological distance *d* from the colony to a location *x* as

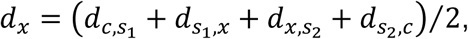

where *d*_*c,s*1_ is the great circle distance from the colony to *s*_l_, the location at which the outward commuting bird crosses the coast; *d*_*s*1,*X*_ is the distance by sea from there to *x*; *d*_*X,s*2_ is the is the distance by sea from *x* to *s*_2_, the inward coast crossing; and *d*_*s2*,*c*_ is the great circle distance from there to the colony (Fig. S2). We calculated *d*_*s1*,*X*_ and *d*_*X,s*2_ on a 3.125 km regular grid using the gdistance package [15]. Location resolution was insufficient to determine where GLS-tracked birds crossed the coast, so we assumed they did so at the median longitudes at which the GPS-tracked birds did so.

**Fig. S2.**
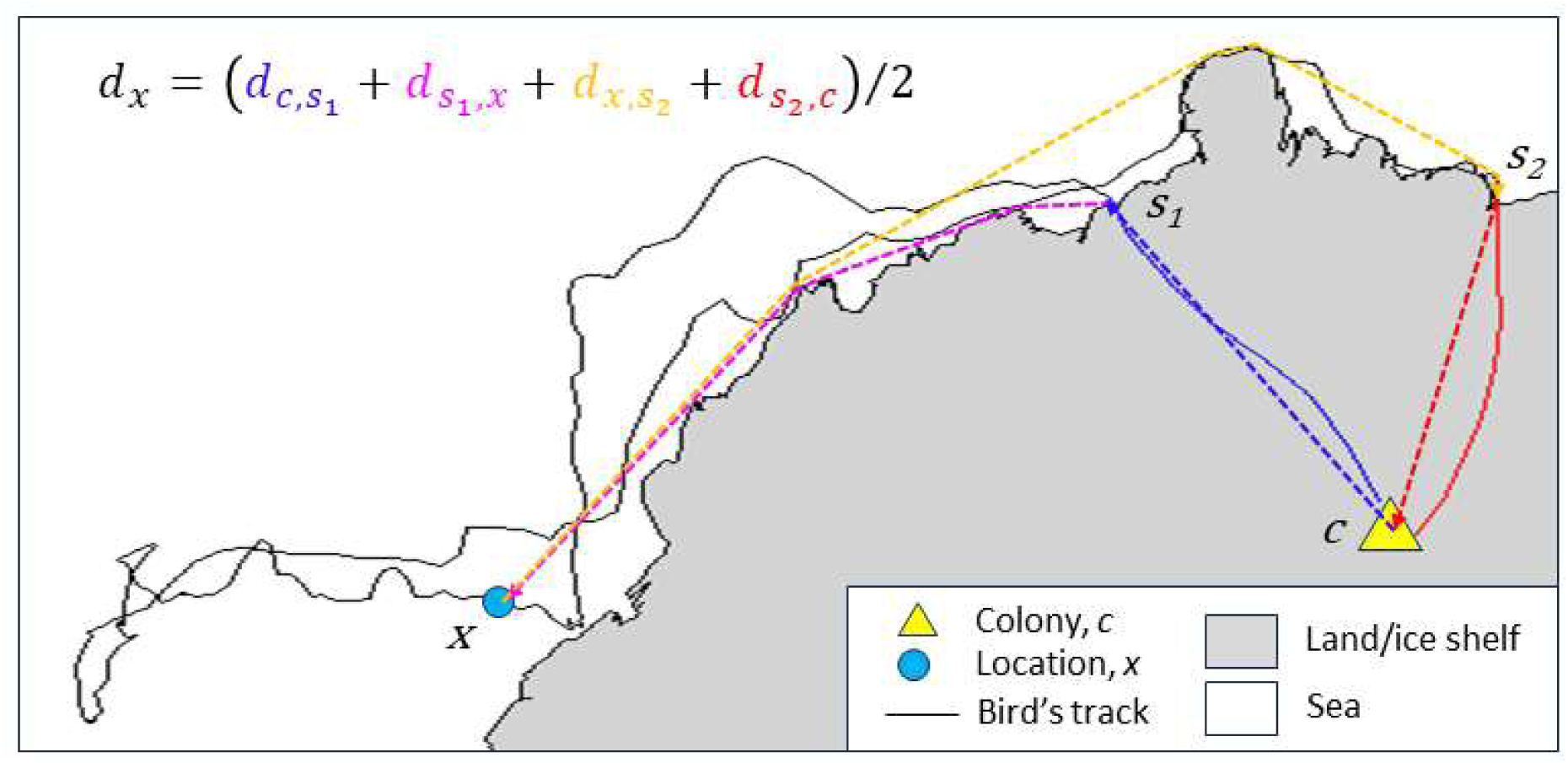
Example of track of a foraging trip made by a snow petrel tracked via GPS from Svarthamaren, Dronning Maud Land during post-brood, illustrating how we defined biological distance, *d* between the colony, *c* and an example location, *x* and passing thought the outward and inward coast crossing locations, *s*_l_ and*s*_2_.

## S2. Supplementary Tables

**Table S1.** Initial and estimated parameter values in Hidden Markov Models of behavioural state of snow petrels GPS-tracked from three colonies in Dronning Maud Land.

| Parameter |  |  | State |  |  |
| --- | --- | --- | --- | --- | --- |
|  |  |  | Resting | Foraging | Travelling |
| Step length<br>(m/s) | mean | initial range | 0.2-1.0 | 3.0-5.0 | 9.0-15.0 |
|  |  | estimate | 0.6 | 4.2 | 8.5 |
|  | sd | initial range | 0.2-1.0 | 3.0-5.0 | 9.0-15.0 |
|  |  | estimate | 0.6 | 2.3 | 3.2 |
| Angle<br>(degrees) | mean | initial | 0.0 | 0.0 | 0.0 |
|  |  | estimate | 0.0 | 0.0 | 0.0 |
|  | concentration | initial range | 0.5-0.8 | 0.5-5.0 | 15.0-25.0 |
|  |  | estimate | 0.7 | 1.4 | 10.5 |

**Table S2.** Mean masses (g) ± sd (range) of snow petrels sampled at the three study colonies in Dronning Maud Land.

| Colony | Females | Males |
| --- | --- | --- |
| Jutulsessen | 232 ± 24 (205 -275; n = 15) | 242 ± 27 (210 -315; n = 16) |
| Svarthamaren | 258 ± 32 (220 -340; n = 15) | 287 ± 40 (230 -370; n = 25) |
| Utsteinen | 257 ± 31 (205 -325; n = 17) | 264 ± 19 (230 -300; n = 14) |

**Table S3.** Linear model of the mass of snow petrels weighed in colonies in Dronning Maud Land vs. sex and colony (models containing interactions between sex and colony (ΔAIC = 1.3), just sex (ΔAIC = 4.6) or just colony (ΔAIC = 19.7) were less parsimonious).

| Covariate | Mass (g) | SE | *t* | *p* |
| --- | --- | --- | --- | --- |
| Jutulsessen, female (intercept) | 228.9 | 6.4 | 35.57 | <0.001 |
| Svarthamaren | 37.5 | 7.5 | 5.03 | <0.001 |
| Utsteinen | 24.3 | 7.9 | 3.08 | <0.001 |
| Male | 16.0 | 6.2 | 2.56 | 0.010 |

**Table S4.** Linear models of median foraging range *d*_50_ of snow petrels tracked via geolocation from three colonies in Dronning Maud Land during pre-breeding (the ten days prior to colony arrival) and the pre-laying exodus vs. breeding season (stages modelled separately).

| Stage | Breeding season | $d_{50}$(km) | SE | *t* | *p* |
| --- | --- | --- | --- | --- | --- |
| Pre-breeding | 2020/21 (intercept) | 1754 | 104 | 16.92 | <0.001 |
|  | 2021/22 | 177 | 147 | 1.20 | 0.235 |
|  | 2022/23 | -103 | 120 | -0.86 | 0.395 |
|  | 2023/24 | -515 | 115 | -4.48 | <0.001 |
| Pre-laying exodus | 2020/21 (intercept) | 1670 | 85 | 19.62 | <0.001 |
|  | 2021/22 | -354 | 120 | -2.94 | 0.005 |
|  | 2022/23 | -242 | 94 | -2.56 | 0.013 |
|  | 2023/24 | -411 | 94 | -4.35 | <0.001 |

## S3. Supplemenatary Figures

**Fig. S3.**
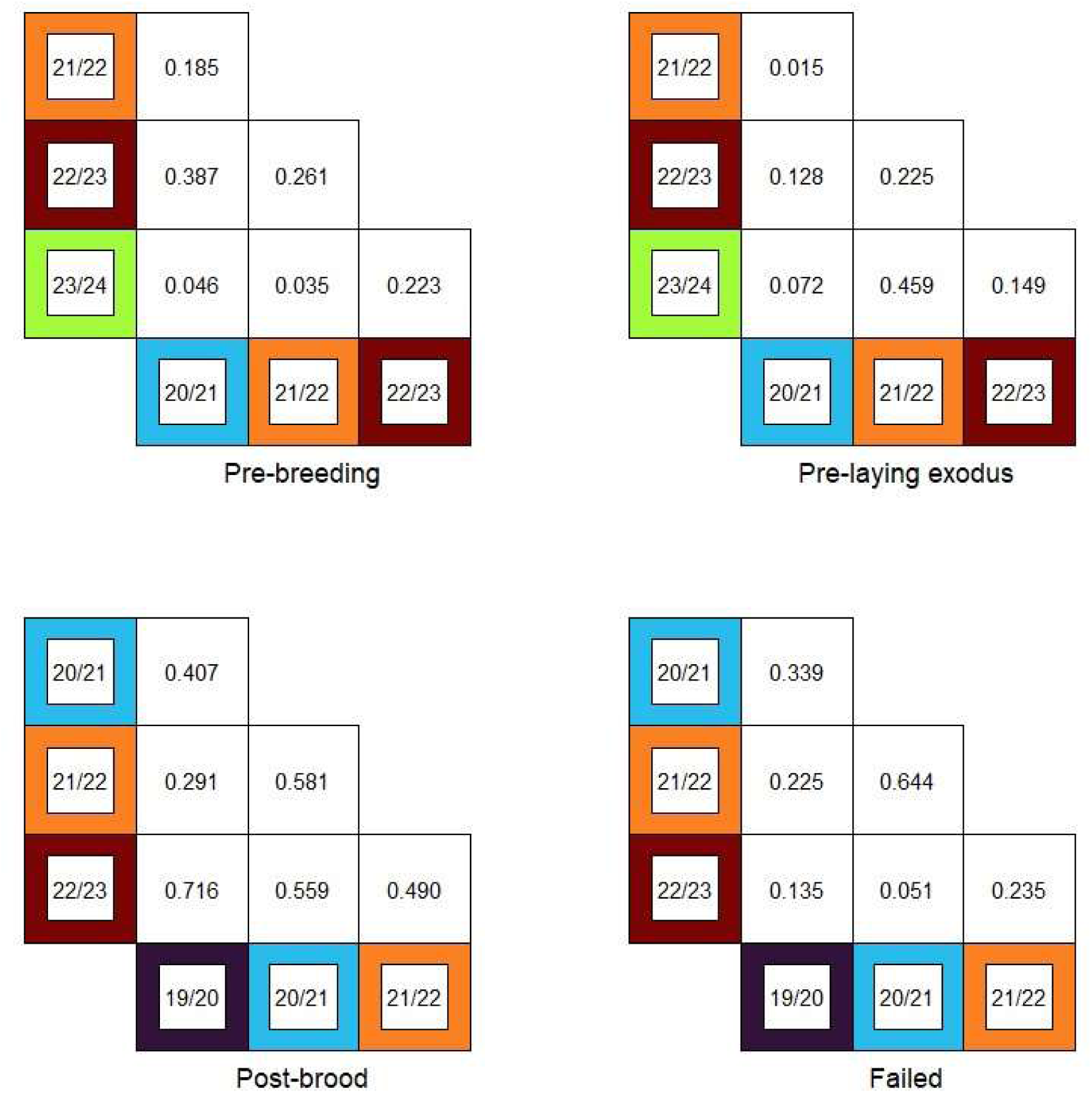
Home Range Overlap Index between core home ranges of birds GLS-tracked from Svarthamaren, Dronning Maud Land during five breeding seasons (colours correspond to Figs. 1 and 2).

**Fig. S4.**
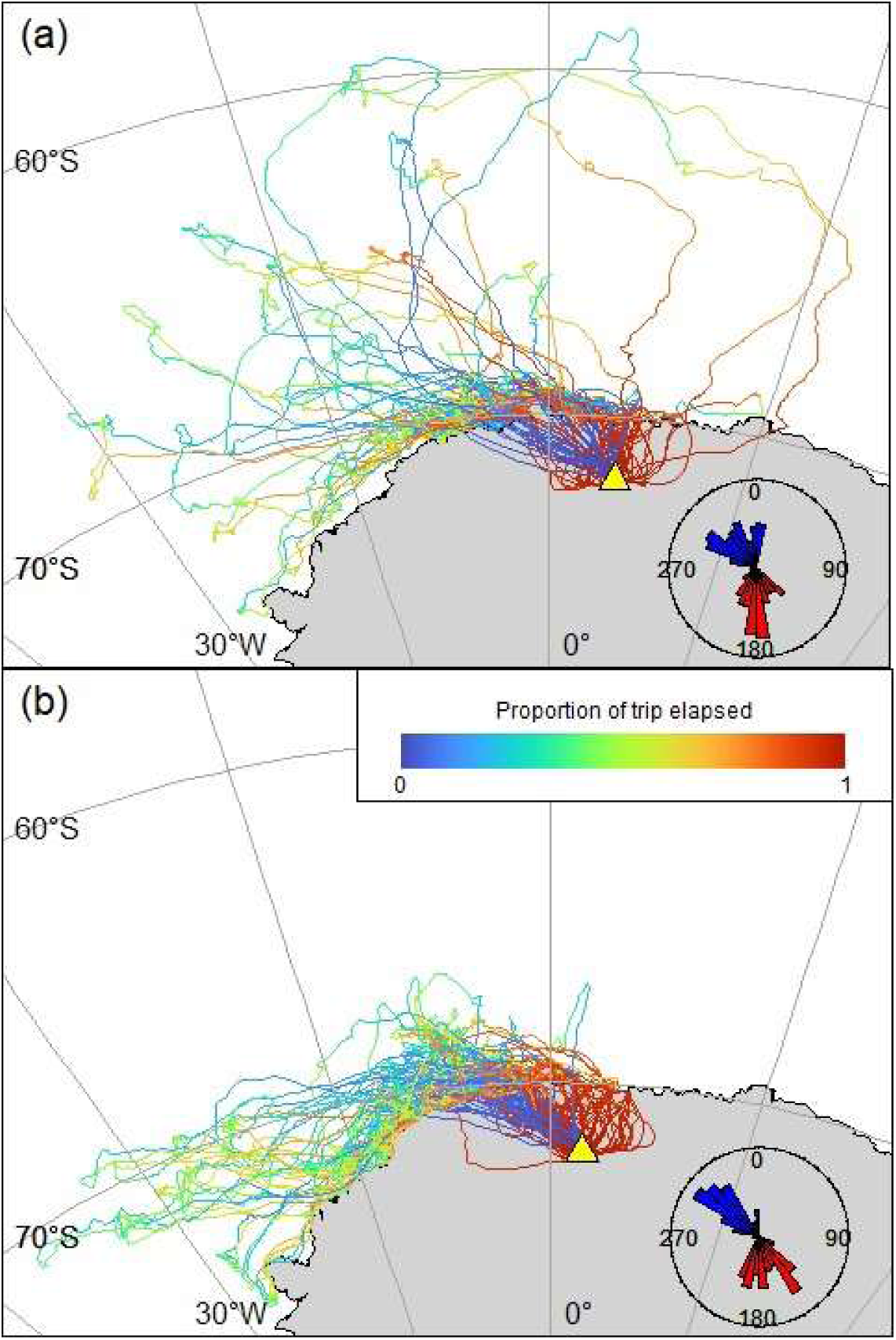
Foraging trips of snow petrels GPS-tracked from Svarthamaren (incubation, brood-guard and post-brood, 2022/23) and Jutlsessen (post-brood, 2024), Dronning Maud Land coloured by proportion of the trip elapsed. Insets show the distributions of mean bird headings on outward (blue) and inward (red) overland commuting legs. Yellow triangles show study colonies.

**Fig. S5.**
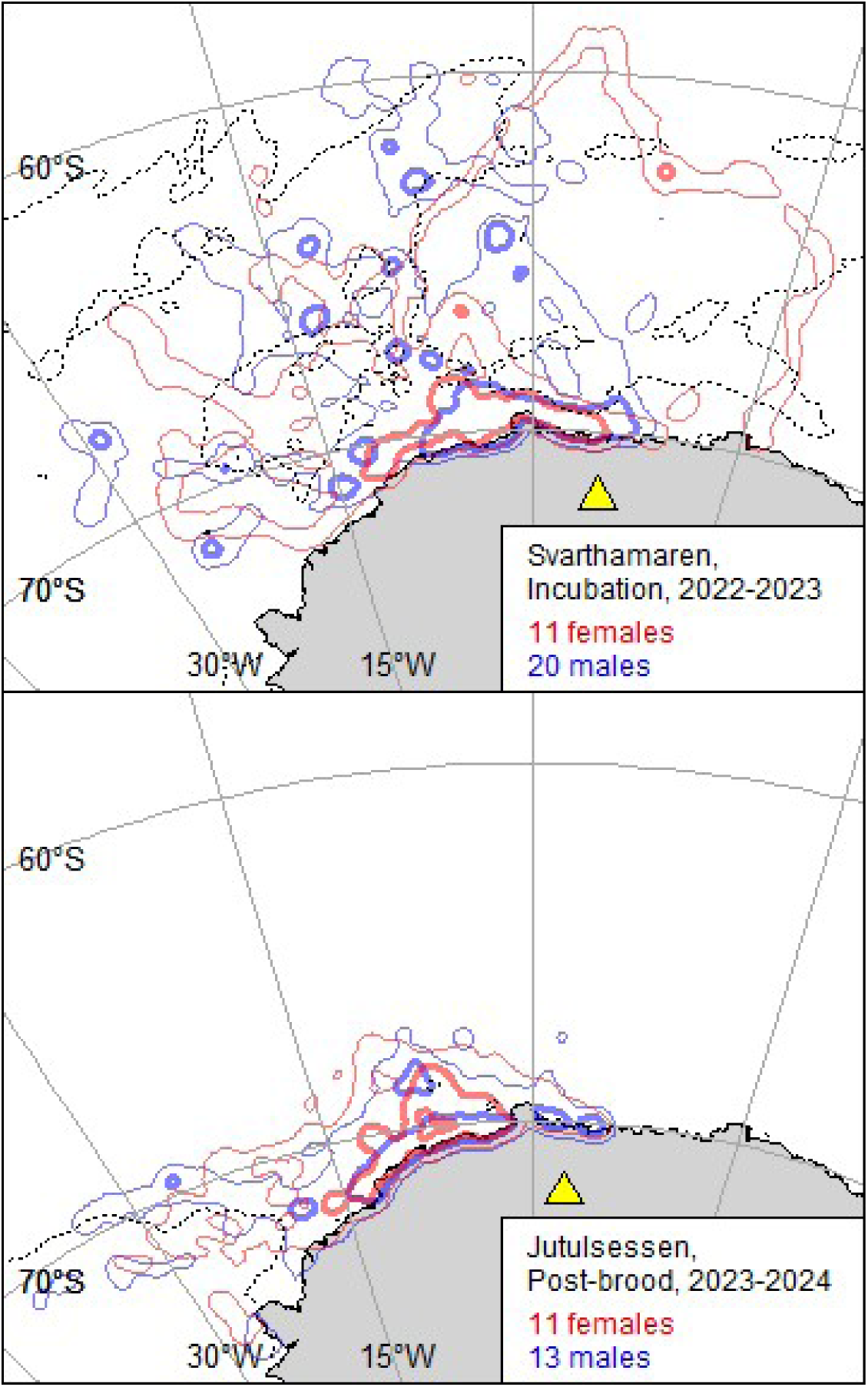
Home ranges of female and male snow petrels GPS-tracked from Svathamaren during incubation and Jutulsessen, Dronning Maud Land during post-brood chick rearing. Thick and thin solid lines indicate 50 and 95 % utilisation volume contours; dashed lines the median ice edge (15% sea-ice concentration) during the corresponding periods; and yellow triangles study colonies.

